# The endocytic recycling pathway is controlled by the ADP-ribosylated GTPase Rab14

**DOI:** 10.1101/2022.11.26.517555

**Authors:** Annunziata Corteggio, Matteo Lo Monte, Laura Schembri, Nina Dathan, Simone Di Paola, Giovanna Grimaldi, Daniela Corda

## Abstract

The GTPase Rab14 is localized at the trans-Golgi network and at the intermediate compartment associated to sorting/recycling endosomes-like structures of the transferrin-recycling pathway: as other Rab family members, it is involved in the regulation of intracellular vesicle trafficking, though its role and functional relationship with effector/endosomal proteins is still incomplete.

We have analysed whether post-translational modifications could affect Rab14 activity: the results obtained define mono-ADP-ribosylation (MARylation) as the yet-unknown Rab14 modification, catalysed by the ADP-ribosyltransferase PARP12, which specifically modifies glutamic acid residues in position 159/162. This modification is essential for the Rab14-dependent endosome progression. Accordingly, recycling of the transferrin receptor is inhibited when MARylation of Rab14 is prevented by PARP12 knocking-down or inhibition, or by overexpression of Rab14 ADP-ribosylation-defective mutant. Under these conditions, Rab14 and transferrin receptors are withheld at the cell periphery at the level of the Rab4-RUFY1-positive sorting endosomes, indicating that the interaction of Rab14 with the dual effectors RUFY and then FIP1c (which specifically binds both Rab11 and Rab14) determines the progression between the Rab4-RUFY- and Rab11-FIP1c-specific vesicles. Therefore Rab14-MARylation determines the sequential binding of this GTPase to RUFY and FIP1c, thus controlling endosome progression (*i*.*e*., transferrin receptors recycling) through the Rab4-, Rab14- and Rab11-specific vesicles. This identifies a Rab14-specific compartment of the recycling pathway and a crucial enzymatic reaction amenable to pharmacological control.

## Introduction

Intracellular membrane trafficking among organelles occurs through vesicular or tubular intermediates that selectively deliver proteins and lipids from one compartment to another ^1^. In mammals, among the different factors that regulate intracellular membrane trafficking, a fundamental role is played by the over 60 members of the Rab GTPase protein family^2, 3^. Rabs are cytosolic proteins that associate with specific organelles when modified (via geranyl-genarylation at the C-terminus); the unique topology of the individual Rab made them reliable markers of the diverse subcellular compartments where they recruit effectors that determine the specificity and directionality of the secretory or endocytic pathways^1, 3-6^. As other GTPases, Rabs act as molecular switches, changing conformations between an active GTP-bound and an inactive GDP-bound state; this GTP/GDP cycle determines the recognition of specific effectors (*e*.*g*., GAP or GEF) involved in the specific Rab activities^7, 8^.

More specifically, Rab4, -5, -7, -9 and -11 are part of the endocytic pathway^9^, which follows two routes: a “degradative endocytic pathway” that brings the cargoes to the lysosomal compartment for degradation, and the “recycling pathway” that brings back the cargoes to the plasma membrane^10^. The “shared” early endocytic pathway is identified and regulated by Rab5; this is followed by either the recycling pathway-governed by Rab4 and Rab11 -or the lysosomal pathway-governed by Rab7 and Rab9^11-14^. In particular, the action of Rab5 in the endolysosomal system characterizes a maturation cascade where Rab5, by recruiting the Rab7-GEF Mon1-Ccz1, activates this Rab favouring the transition of the endosomes Rab5-specific into Rab7-specific, that then finally fuse with lysosomes^15^. This well-defined Rab cascade, determines the maturation of endosomes towards lysosomes bringing the cargoes to the final station^7-9, 16^.

At steady-state Rab14 localizes at the trans-Golgi network (TGN) and at recycling endosomes, intervening in both secretory and recycling endocytic pathways^17-19^. The recycling pathway can occur through a direct route from the sorting endosomes to the plasma membrane (the so-called “fast-recycling pathway”) or, indirectly, *via* the more specialized endosomal recycling compartment (ERC), that involves a transit at the perinuclear area, before returning to the plasma membrane (the “slow-recycling pathway”)^1, 9, 20, 21^. Rab14 has been associated to the Rabs regulating the endosomal recycling pathways, at the Rab11-specific endosomes, at the transition step between sorting endosomes and perinuclear ERC^22^, although its specific mechanism of action has not been identified^19, 23^.

Nevertheless, Rab14 action appears to be relevant in diverse diseases^24^: Rab14 overexpression has been associated to carcinogenesis in different tumors and its function as an oncogene hypothesized^25-28^ Also, Rab14 has been involved in viral infection/internalization and recycling^29-31^. These recent observations raise the interest on Rab14 not only for its role in membrane transport, but as a potential target for drug development^24, 28, 32^.

Beside the GTPase activity, an additional regulation of Rabs functions is related to their post-translational modifications (PTMs). So far, phosphorylation, and also dephosphorylation, have been shown to interfere with Rab functions, as in the case of Rab1^33^ or Rab7^34, 35^. Similarly, adenylylation (attachment of adenosine monophosphate, AMP, known also as AMPylation) of human Rab1b by the bacterial enzyme DrrA/SidM from *Legionella pneumophila* blocks the access of GTPase activating proteins, thereby making Rab1b constitutively active^36, 37^.

Among the PTMs potentially regulating proteins involved in membrane traffic, we have been investigating the role of mono-ADP-ribosylation (MARylation).

MARylation is a reversible PTM consisting in the enzymatic transfer of ADP-ribose from NAD^+^ to specific amino-acids of target proteins, with the simultaneous release of nicotinamide^38^. Originally identified in bacteria as a pathogenic mechanism to infect host cells^39^, it has been later described also as an endogenous PTM to regulate protein functions^40-43^. PARP enzymes are ADP-ribosyltransferases able to catalyse this reaction intracellularly^44^, both under stress and steady-state conditions^45, 46^. The human PARP family is composed of 17 members, characterized by a conserved catalytic domain and additional diversified domains, mediating DNA/RNA-binding, ADP-ribose-recognition, protein-protein interactions, thus determining specific functions and subcellular localizations^47^.

Our lab has mainly contributed to the elucidation of the endogenous role of MARylation^48, 49^ also reporting that the PARP12-dependent MARylation affects the function of the Golgi complex and hence intracellular membrane traffic^50^. This occurs under stress conditions, when PARP12 is sequestered to stress granules thus impeding TGN to plasma membrane traffic^45^. In addition, this enzyme endogenously MARylates Golgin-97 thus modulating the traffic of E-cadherin toward the PM^46^. Other PARPs may affect diverse cellular functions: this is the case of PARP10-mediated MARylation of the Aurora-A kinase that is required during the G2/M transition of the cell cycle^51^. Altogether these reports contribute to the notion of a functional role of MARylation in diverse cellular events.

Based on the above, we have planned to further investigate the role of PARP12 and dependent MARylation in the control of intracellular membrane traffic by first identifying specific substrates of this reaction, then selecting those specifically involved in traffic steps, and finally analyzing the role of this PTM in their functions.

We report that Rab14 is a specific, direct substrate of PARP12-dependent MARylation and that through this PTM PARP12 controls the recycling pathway of transferrin receptor by ensuring its efficient transition from early/sorting endosomes to perinuclear recycling endosomes.

Specifically, we demonstrate that PARP12 MARylates Rab14 at specific residues and regulates its compartmentalization and activity, thus contributing to critical features for recycling endosomal progression, a physiological event whose defects have been involved in a wide range of diseases, including cancer and neurodegenerative disorders^52, 53^.

## Results

### Rab14 is a substrate of PARP12-catalyzed MARylation

PARP12 plays a role in the exocytic pathway of basolateral cargoes^46^. To further investigate its function in membrane trafficking, we sought to identify PARP12-specific substrates by applying the Af1521 *macro* domain-based pull-down assay (a domain that specifically recognizes ADP-ribose bound to proteins; for brevity Af1521 assay) coupled to mass spectrometry (MS) analysis (see methods and^54^). To this end, a PARP12-enriched membrane fraction of HeLa cells overexpressing PARP12 was used to perform an *in vitro* ADP-ribosylation assay, followed by *macro*-domain pull-down. HeLa cells transfected with empty vector alone were used as control. Af1521-bound proteins obtained in both conditions were subsequently identified by MS analysis (see Methods) and PARP12 specific substrates clustered according to their functions into five major subgroups: proteins involved in RNA processing, DNA-binding proteins, members of the small-GTPase family, mitochondrial proteins and plasma-membrane receptors (Suppl. Fig. 1). Of note and in accordance with our previous data^46^, some of the identified substrates were involved in intracellular trafficking, thus we first focused our studies on one member of this group, the small GTPase Rab14.

To validate Rab14 as a PARP12 substrate, *in vitro* MARylation assays were performed using recombinant-purified, GST-tagged PARP12 catalytic fragment and His-Rab14 in the presence of ^32^P-NAD^+^ (see methods): this resulted in the Rab14 modification (Fig.1a); instead, no modification was observed when tankyrase -an enzyme of the PARP family localized at the *trans* Golgi-was used in *in vitro* ADP-ribosylation assays (Suppl. Fig. 2a). Accordingly, we could conclude that Rab14 is a specific, direct PARP12 substrate.

**Fig. 1:**
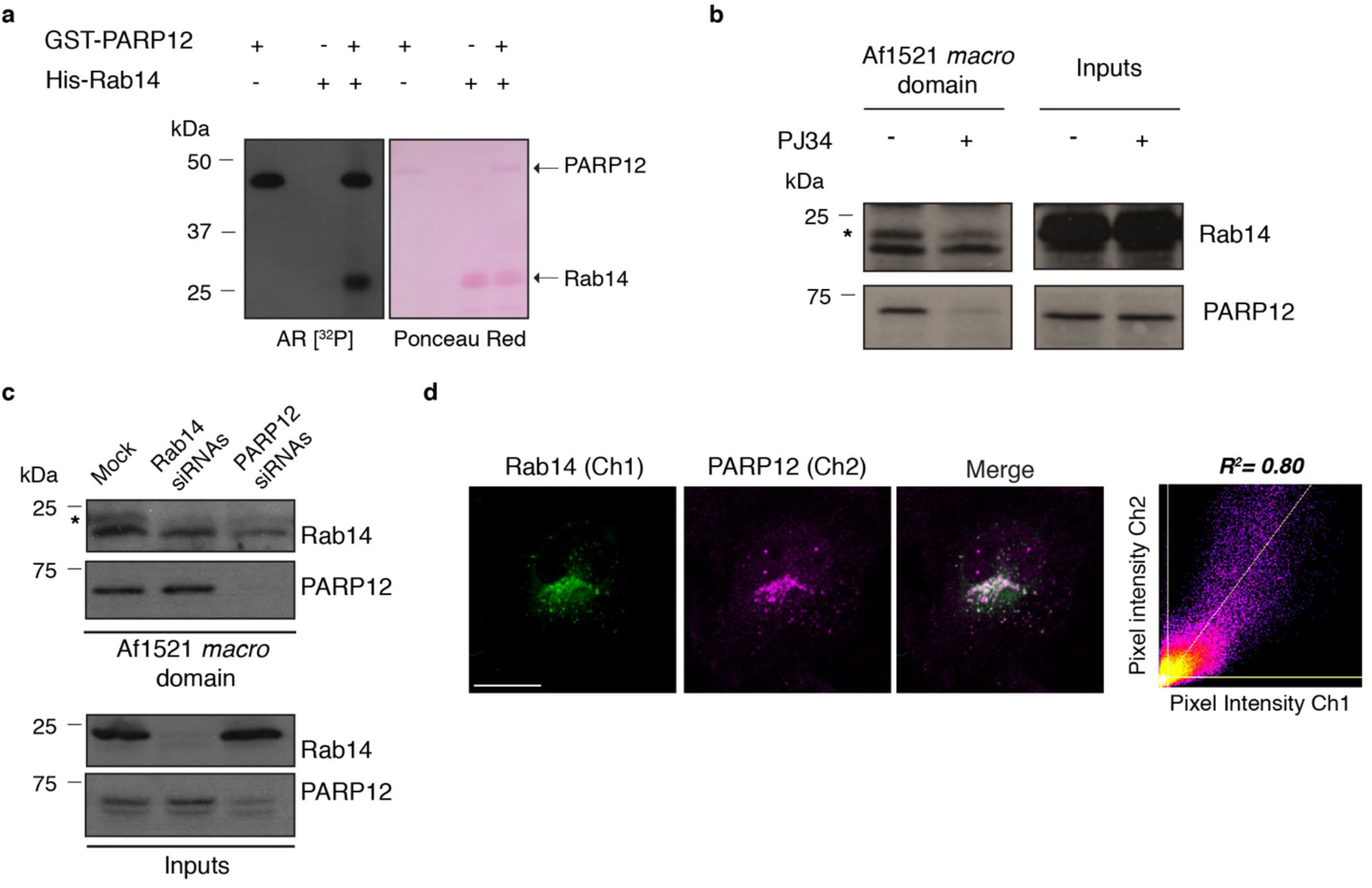
Rab14 is a PARP12 substrate. **(a)** *In vitro* ADP-ribosylation assay using GST-tagged purified PARP12 catalytic fragment and His-tagged purified Rab14, in presence of 4 μCi [^32^P]-NAD^+^ and 30 μM total NAD^+^. The incorporated [^32^P]-ADP-ribose was detected by autoradiography (AR [^32^P]). Ponceau S staining is shown for total levels of PARP12 (upper bands) and Rab14 (lower bands). **(b, c)** Af1521 *macro* domain based pull-down assay of total cell lysates obtained from (b) HeLa cells treated or not with the broad PARP inhibitor PJ34 (50 μM, 2 h) or (c) HeLa cells depleted of Rab14 (Rab14 siRNAs), PARP12 (PARP12 siRNAs) or not (Mock), showing MARylated Rab14 (asterisks). MARylation of PARP12 was detected as internal control. **(d)** Representative confocal images of HeLa cells at steady state, fixed and labeled for endogenous Rab14 (green, Ch1) and PARP12 (magenta, Ch2). Merged signals are also shown. Scatter plot shows the correlation analysis of pixel fluorescence intensities for Rab14 and PARP12 signals. R2 value is indicated. Scale bar, 10 μm.

Rab14 MARylation was also detected in total cell lysates using the Af1521 assay (Fig. 1b, c). Since PARP12 also undergoes auto-modification^45, 55^, its binding to the Af1521 *macro*-domain was considered an appropriate internal control. HeLa cells treated with the broad PARP inhibitor PJ34 (50 μM, 2 h) or transiently depleted of PARP12 expression (by siRNAs) showed a pronounced reduction (respectively of about 45% and 60%, compared to control cells) in endogenous Rab14 binding to the Af1521 *macro*-domain, indicating that endogenous PARP12 specifically MARylates Rab14 in intact cells.

The assay was also performed on total lysates obtained from cells depleted of Rab14 (Rab14 siRNAs) and was used as control of specificity for the Rab14 antibody.

Rab14 binding to the Af1521 *macro*-domain was not affected by treatment with the tankyrase specific inhibitor IWR1 (Suppl. Fig. 2b), in line with the data described above and with the specificity of the PARP12-dependent modification.

Overall, these data demonstrate that MARylation is the first-reported Rab14 post-translational modification, specifically catalyzed by PARP12.

Importantly, Rab14 and PARP12 were localized in the same subcellular compartment at the perinuclear area and cytoplasmic vesicles, as indicated by their co-localization evaluated with specific antibodies (Fig. 1d), further suggesting their potential cooperation in the regulation of traffic events (see below).

### MARylation regulates the intracellular localization of Rab14

The subcellular localization and distribution of Rab proteins is crucial in determining specificity and directionality of membrane traffic. Specifically, Rab14 has been described to localize and act on an intermediate compartment along the transferrin-recycling pathway, prior to Rab11 and after Rab5 and Rab4, where it is specifically involved in the transport of the transferrin receptor from early/sorting endosomes towards perinuclear recycling endosomes, Rab11-positive^22^.

To explore the impact that MARylation might have on Rab14 function, we firstly investigated whether lack of this PTM affected Rab14 localization. HeLa cells were treated with the PARP inhibitor PJ34 (50 μM, 2 h) and the distribution of either endogenous Rab14 or over-expressed EGFP-tagged Rab14 analyzed by immunofluorescence, by using Rab14 antibody or following the EGFP signal (Fig. 2a). At steady-state, Rab14 was detected throughout the cell in small, sorting/recycling endosomes-like vesicles^22, 56^ with a major concentration in the perinuclear area, as previously described^17^. Acute treatment with PJ34 altered Rab14 localization, which mostly accumulated in vesicles dispersed at the cell periphery (Fig. 2a). These Rab14-positive vesicles were analyzed for their co-localization degree with markers of early and recycling endosomes or lysosomes or Golgi complex membranes (see methods). At steady state, Rab14 mainly localized at the ERC and, for a minor fraction, in early/sorting endosomes, positive for both EEA1 and TfR. Inhibition of PARP-mediated ADP-ribosylation resulted in an increased co-localization of Rab14 with both endosomal markers EEA1 and TfR (Fig. 2 b-e), while no colocalization was observed with the lysosomal marker LAMP1 (Suppl. Fig. 3a) or with a marker of the Golgi apparatus (Golgin-97, Suppl. Fig. 3b). Of note, TfR itself was concomitantly partially shifted from the TGN toward a peripheral localization after PJ34 treatment, mimicking the Rab14 distribution under identical conditions (Fig. 2c, g). Treatment with the tankyrase inhibitor IWR1 did not modify Rab14 localization (Suppl. Fig. 2c); on the same line of evidence, tankyrase was not able to catalyze the ADP-ribosylation of Rab14 from cell lysates (Suppl. Fig. 2b).

**Fig. 2:**
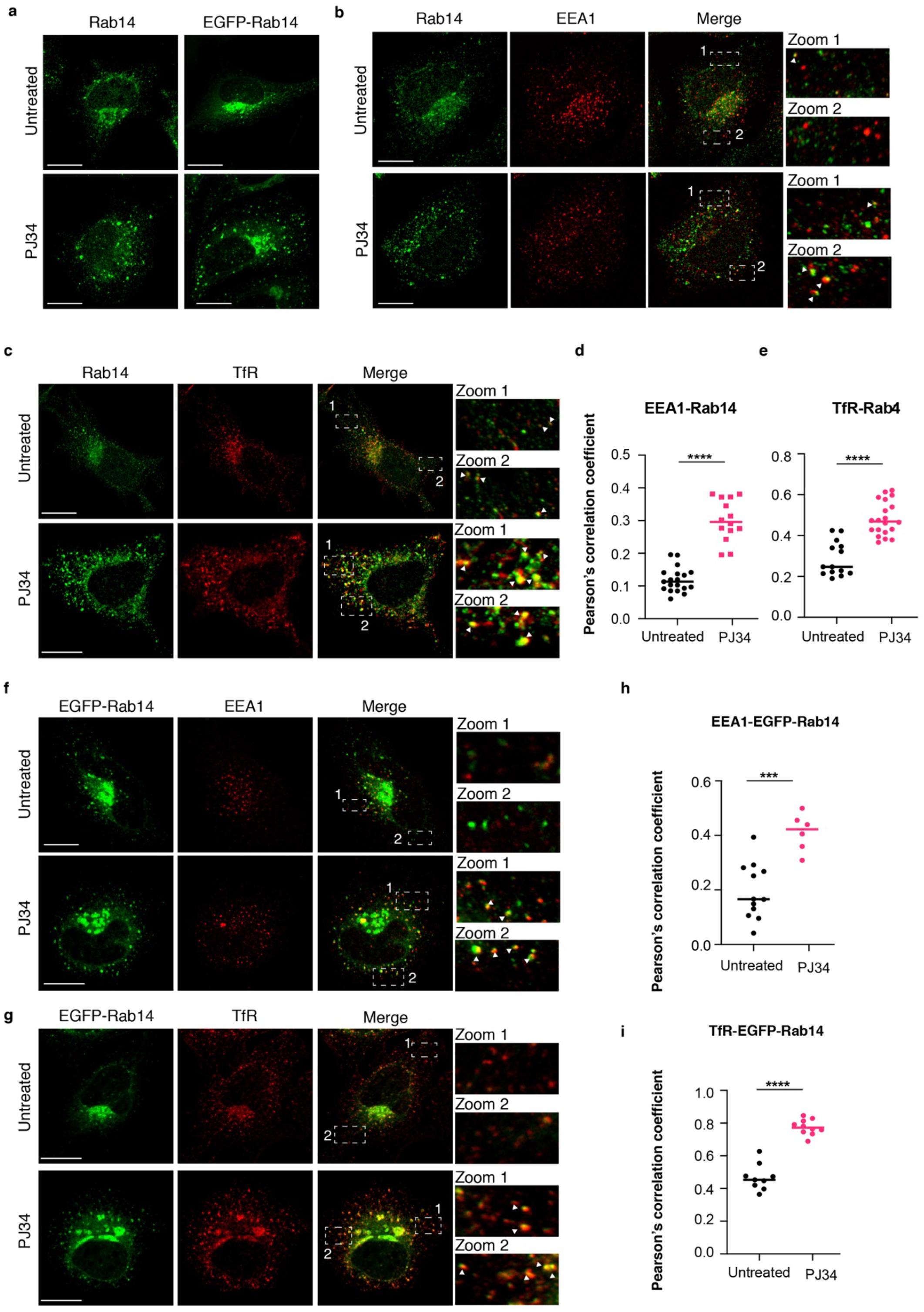
Effect of PJ34 treatment on Rab14 localization. **(a)** HeLa cells transfected or not with EGFP-Rab14 were treated or not with PJ34 (50 μM, 2 h), fixed and labeled with anti-Rab14 antibody. **(b, c, f, g)** Effect of PJ34 treatment (50 μM, 2 h) on localization of endogenous or overexpressed Rab14 (Rab14-eGFP), as indicated; HeLa cells were fixed and labeled with antibodies against (b) Rab14 and EEA1, (c) Rab14 and transferrin receptor (TfR), or (f) EEA1 and (g) TfR alone. Representative confocal microscopy images are reported. Merged signals are also shown. **(d, e, h, i)** Co-localization analysis was performed using Pearson’s correlation coefficient, as indicated. Scale bars, 10 μm.

Together, these data demonstrate that lack of MARylation induces Rab14 accumulation in enlarged early/sorting endosomal structures (Fig. 2), an event that may cause a general block of the entire endosomal progression.

To directly evaluate PARP12 involvement in determining this altered phenotype, Rab14 localization was analyzed in PARP12 knocked-down cells. While in control cells (Mock) Rab14 mainly accumulated in the perinuclear area, in the absence of PARP12 it was mainly present at the cell periphery in large spots positive for EEA1 and TfR (Fig. 3a, b), as also observed upon PJ34 treatment (Fig. 2). In parallel, we evaluated the effects of both PJ34 treatment and PARP12 depletion on the localization of Rab4, a member of the same Rab protein subfamily involved in the early steps (sorting) of the TfR endocytic pathway. None of these treatments affected Rab4 localization (Suppl. Fig. 4a, b), indicating that the PARP12-mediated catalytic activity is specifically regulating Rab14 localization, clearly acting downstream of and independently from Rab4 activity.

**Fig. 3:**
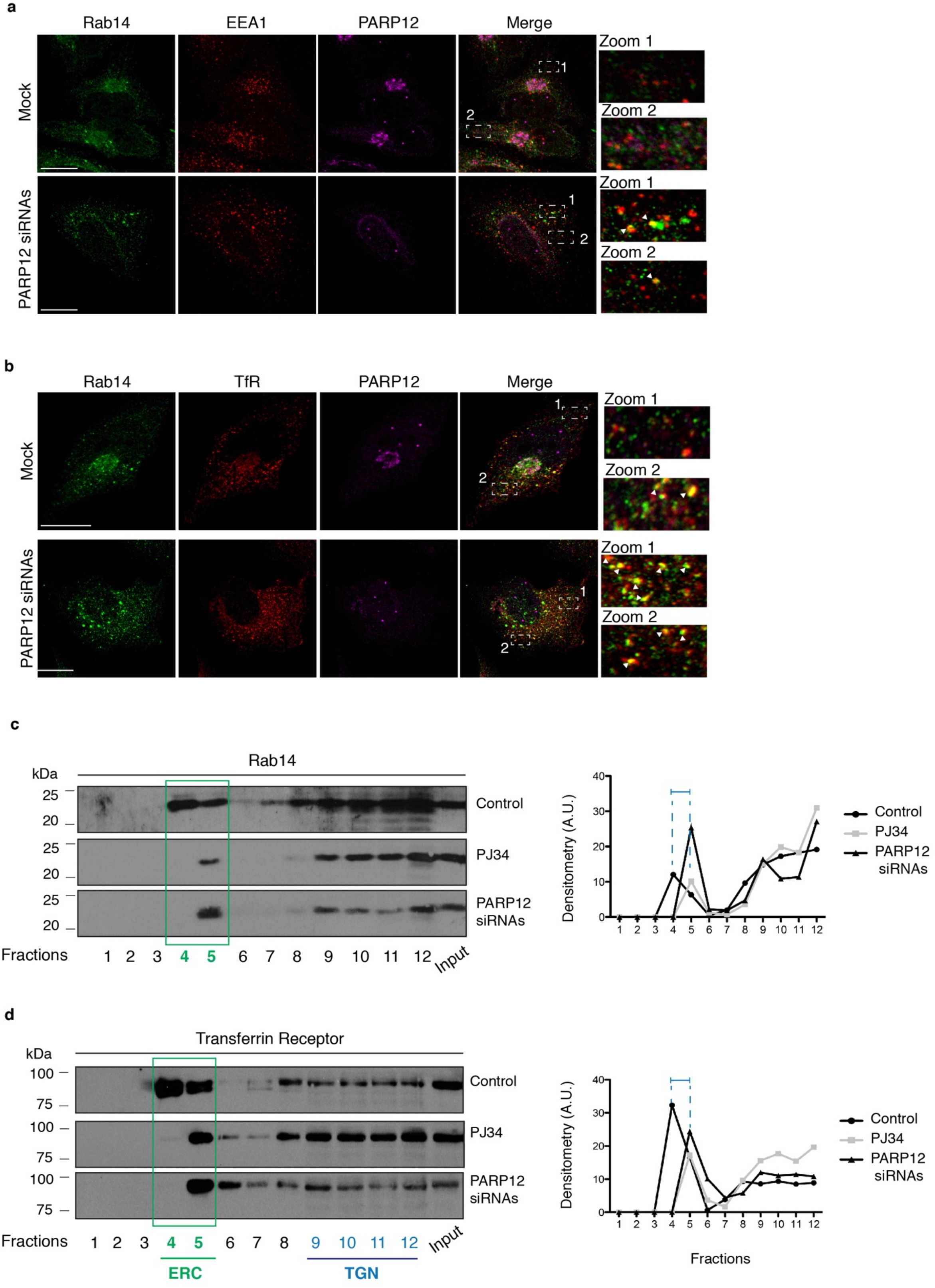
Effect of PARP12 depletion on Rab14 localization. **(a, b)** Representative confocal microscopy images of endogenous Rab14 localization upon PARP12 depletion using PARP12 siRNAs. Mock samples used as control. After 72 h, cells were fixed and stained with antibodies against Rab14 (green), EEA1 or TfR (red) and PARP12 (magenta). Zoom 1 and 2: enlarged views of merged signals. Scale bars, 10 μm. **(c)** Representative detergent-free isolation of buoyant cholesterol-rich endomembrane fractions (ERC/TGN enriched-fractions) of Hela cells untreated (Control), treated with PJ34 (50 μM, 2 h) or transfected with PARP12 siRNAs. Equal volumes of all subcellular fractions and equal amount of total protein lysates before fractionation (input) were analyzed by western blotting for Rab14 and Transferrin receptor and each fraction quantified as percentage of the sum of protein abundance.

Rab14 localization depends on an association/dissociation cycle to/from membranes that is superimposed to the GTP/GDP cycle^57^. Our data show a change in Rab14 localization when MARylation is hampered (Figs 2, 3). On this basis, we analyzed Rab14 cytosol/membrane partitioning (see Methods) and show that it is mainly associated to the total membrane fraction and, to a lesser extent, to the cytosolic fraction. This distribution was not altered by PARP12 depletion or by PJ34 treatment (Suppl. Fig. 4c), indicating that PARP12-mediated MARylation is not affecting Rab14 association to the total membrane pool, rather it affects Rab14-effector interactome along the recycling pathway (see Figs 2,3, Suppl. Fig. 4).

We also evaluated Rab14 distribution within the subcellular endomembrane fractions obtained by differential centrifugation followed by sucrose-gradient separation (see Methods). Isolation of pericentrosomal ERC and post TGN-vesicles (buoyant cholesterol-rich endomembranes, fractions 4-5) showed that Rab14 was present in these fractions, as indicated by TfR labelling (Fig. 3 c, d, lanes 4-5); the amount of Rab14 present in the ERC fractions decreased (Rab14 was present only in fraction 5) after PJ34 treatment or when PARP12 was depleted (Fig. 3c); a similar trend was also observed for TfR (Fig 3d).

Collectively, our data indicate that the subcellular compartmentalization of Rab14 is controlled by PARP12-mediated MARylation: inhibition of this PTM impairs Rab14 recruitment to perinuclear-recycling endosomes close to the TGN region, while causing a prolonged peripheral endosomal localization (Figs 2, 3).

### MARylated Rab14 is required for efficient recycling of transferrin receptor

As detailed above, inhibition of PARP12-dependent MARylation also resulted in a defect in the TfR distribution (Figs 2, 3), supporting the concept that MARylation has a role on the entire endosomal recycling process mediated by Rab14.

Based on the above, to assess this point, we analyzed the endocytic pathway of the TfR, as specific cargo regulated by Rab14^22^. Fluorescently-labeled transferrin was used to track the TfR recycling pathway (see Methods), in both control cells and in cells defective of PARP12-mediated MARylation (obtained by pre-treatment with either the PJ34 inhibitor or specific siRNAs as reported above; see methods).

Upon binding to its receptor, transferrin is endocytosed and trafficked to peripheral early endosomes, to then move to recycling endosomes, concentrated in the perinuclear ERC, and finally be delivered back to the cell surface^58^. HeLa cells were allowed to internalize Alexa-fluor-633-labeled transferrin (Alexa-633 transferrin) at 37 °C and, as expected, after 30 min of continuous internalization, control cells showed a bulk of transferrin bound to its receptor targeted to the perinuclear ERC, a localization coincident with that of Rab14 (Fig. 4a, Suppl. Fig 5a, upper panels). Differently, cells treated with PJ34 or which were PARP12-depleted showed the internalized transferrin accumulated in peripheral enlarged endosomes, positive for Rab14 and RUFY1, a marker of early/sorting endosomes (Fig. 4 a, Suppl. Fig 5a, lower panels).

**Fig. 4:**
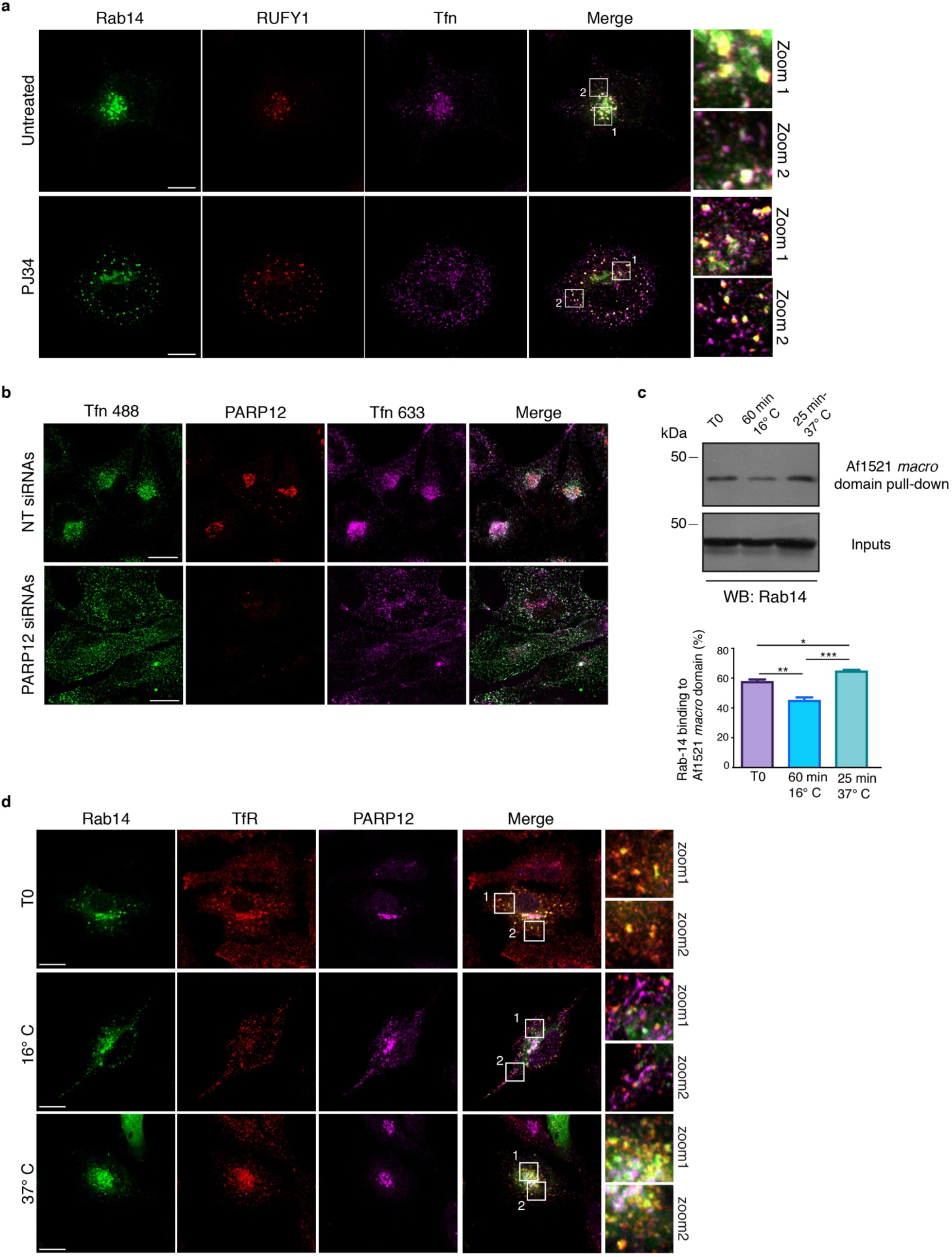
PARP12-mediated MARylation impairs Rab14-driven endocytic recycling traffic. **(a)** Representative confocal microscopy images of Alexa-633-Transferrin (Tfn) uptake in HeLa cells treated or not with PJ34 treatment (50 μM, 2 h), upon EGFP-tagged Rab14 overexpression. After treatments, cells were processed for immunofluorescence using a RUFY1antibody (red). Zoom 1 and 2: enlarged views of merged signals. **(b)** Representative confocal microscopy images of double-transferrin internalization assay performed in HeLa cells transfected first with non-targeting siRNAs or PARP12 siRNAs and then transfected with EGFP-tagged Rab14. After the double pulse of transferrin load (Alexa-488/633-Transferrins), cells were fixed and stained using a PARP12 antibody (red). Merged signals are also shown. **(c)** Rab14 MARylation analysis by Af1521 *macro* domain pull-down assay on total lysates from HeLa cells transfected with EGFP-tagged Rab14, and untreated (T0) or incubated with biotinylated-transferrin (50 μg/ml) for 1 h at 16 °C or 25 min at 37° C (as indicated). Quantifications of MARylated Rab14 are reported in the graphs. Data represent the mean ± S.D. (N = 3 independent experiments; two-sided one-sample t-test; *P <0,05, **P <0,005, ***P <0,001). **(d)** Representative confocal microscopy images of HeLa cells transfected with EGFP-tagged Rab14 and incubated or not (T0) with biotinylated-transferrin (50 μg/ml) for 1 h at 16 °C or 25 min at 37 °C, as indicated. At the end of incubation, cells were fixed and processed for immunofluorescence using TfR (red) and PARP12 (magenta) antibodies. Merged signals are also shown. Zoom 1 and 2: enlarged views of merged signals. Scale bars, 10 μm.

RUFY1 is a bivalent-effector molecule with the ability to bind both Rab4 and Rab14^23^, possibly connecting the two Rabs and thus coordinating respectively the entry and exit of transferrin from early/sorting endosomes. This knowledge, combined with our data showing an accumulation of transferrin in RUFY1-positive endosomes in the absence of MARylation, supports the conclusion that, due to MARylation-defective conditions, transferrin bound to its receptor is trapped at the exit of early/sorting endosomes thus failing to reach the perinuclear ERC. This observation supports the concept that MARylated Rab14 plays a role in the endosomal progression.

To directly evaluate the dynamic behavior of Rab14 during endosomal traffic, we performed live-cell time-lapse imaging experiments of transferrin-uptake in HeLa cells expressing EGFP-tagged Rab14. Cells were incubated in an uptake medium containing Alexa-546 transferrin at 37 °C and immediately imaged for 60 min (see methods, and ^22^). Under conditions of continuous internalization, in control cells, the bulk of labeled transferrin bound to its receptor was present in small vesicles, Rab14-positive, that were continuously moving and targeted to perinuclear ERC (see Suppl. Video 1). Conversely, in PARP12 knocked-down cells, labeled transferrin was clustered in enlarged peripheral Rab14-positive endosomes, that remained static and failed to be targeted to pericentrosomal ERC (see Suppl. Video 2). These set of data demonstrates that PARP12 drives TfR transport from early/sorting endosomes to ERC, while not affecting initial steps of the pathway (*i*.*e*., internalization and early endosomal localization). To further support this finding, cells were double-labeled with two consecutive pulses of distinctly labeled transferrin species *i*.*e*., Alexa-633 and -488 transferrin (see methods and ^59^). Then Hela cells, depleted or not of PARP12, were allowed to internalize first Alexa-633 transferrin for 60 min and then, Alexa-488 transferrin for additional 15 min. In control cells, both transferrins co-localized at perinuclear ERC, as indicated by merged signals deriving from 633- and 488-labeled transferrins (Fig. 4b, upper panel), indicating that the endocytic pathway was fully functional. Differently, in cells depleted of PARP12, the two transferrins were accumulated in separated peripheral enlarged endosomes, which failed to reach the pericentrosomal ERC (Fig. 4b, lower panel), demonstrating that the first steps of the recycling pathway are not defective upon PARP12 depletion.

Collectively, these results indicate that MARylated Rab14 is required for a proper progression of endosomes from the early/sorting station to the perinuclear ERC, thus constituting a new regulatory step along the slow-recycling endocytic pathway (see model Fig. 7).

### MARylation of Rab14 is modulated during transferrin receptor trafficking

The data obtained so far support the conclusion that MARylation of Rab14 is functional to the transferrin trafficking between early/sorting endosomes and recycling endosomes, as indicated by the defective Rab14 localization and defective-transferrin transport caused by the impairment of MARylation. To further investigate this aspect, MARylation of Rab14 was monitored in intact cells under conditions known to allow the traffic of endocytosed transferrin either into early endosomes or to perinuclear ERC^59^. To synchronize the arrival of transferrin into early endosomes, Hela cells expressing EGFP-tagged Rab14 were allowed to internalize biotinylated transferrin for 60 min at 16 °C, a temperature that is permissive for transferrin internalization but not for its exit from the early endosomes. Similarly, to synchronize transferrin into recycling endosomes, HeLa cells expressing EGFP-tagged Rab14 were allowed to internalize biotinylated transferrin for 25 min at 37 °C, a condition that allows the targeting of transferrin to perinuclear ERC. The Af1521 assay was performed under these conditions: Rab14 binding to Af1521 *macro* domain increased by about 35% when transferrin was continuously internalized at 37 °C (*i*.*e*. transferrin at perinuclear ERC), compared to cells allowed to internalize transferrin at 16 °C (*i*.*e*. transferrin at early endosomes, Fig. 4c), indicating that Rab14-endogenous MARylation is triggered by the endocytic stimuli.

Finally, the localizations of both PARP12 and Rab14 were analyzed during the transferrin-uptake assay under the conditions reported above. Upon Alexa-546-transferrin internalization at 16 °C, PARP12 translocated from the TGN to newly-formed early endosomes, positive for Rab14 and transferrin (Fig. 4d). When transferrin was continuously internalized at 37 °C, PARP12 re-located at the TGN, thus suggesting that PARP12 was able to translocate under endocytic stimuli (Fig. 4d). In addition, PARP12 localization was monitored in cells treated with the endosomal recycling inhibitor primaquine, a drug which accumulates in endosomes and inhibits membrane recycling^60^. Similar to the 16 °C temperature block, in the presence of primaquine, PARP12 accumulated in endosomes positive for Rab14 and transferrin (Suppl. Fig. 5b). This treatment did not alter the localization of other Golgi localized proteins such as Golph3 (Suppl. Fig. 5c), indicating that the Golgi complex organization was not affected during this treatment.

The conclusion that can be drawn from the above data is that Rab14 MARylation occurs during transferrin trafficking: upon endocytic stimuli, PARP12 translocates from the Golgi complex to the newly-formed Rab14-positive endosomes, where it modifies and consequently regulates Rab14 functions required for endosomal progression.

### Identification and validation of Rab14 MARylated residues

Having defined Rab14 as a substrate of PARP12-mediated MARylation, we set-out to identify its modified residue(s). To this end we applied the bioinformatic tool ADPredict^61^ designed to predict the acidic residues (aspartic and glutamic acids, D and E) most prone to MARylation (see Methods). Four residues were identified as reliable candidates for ADP-ribosylation, namely E159, E162, E187 and E205 (Table S1). Taking advantage of the close proximity of the first two candidates, E159 and E162, a double point mutant approach was preferred for a first round of testing. We generated a Rab14 multiple point mutant, obtained by substituting E159 and E162 with the relative amide (Glutamine, Q instead of glutamic acid, E). This double-point mutant was first analysed for its ADP-ribosylation levels in *in vitro* assays, using GST-tagged PARP12-catalytic fragment, ^32^P-NAD^+^ and His-Rab14 (wild-type or E159Q-E162Q mutant) for different times. The Rab14 mutant showed a strong reduction in MARylation compared to wild-type Rab14, detectable at all time points, most pronounced at 5 min (about 70%, Fig. 5a, b). The Rab14 mutant modification was also tested in intact cells. To this end, the Af1521 assay was performed on cell lysates of Hela cells transiently overexpressing EGFP-tagged Rab14 wild-type or its E159Q-E162Q mutant and the pool of Af1521-bound Rab14 detected using a Rab14 antibody. This mutant showed a 50% reduction in Af1521 *macro*-domain binding (Fig. 5c, d), thus validating the E159Q-E162Q double-point mutant as defective in MARylation.

**Figure 5:**
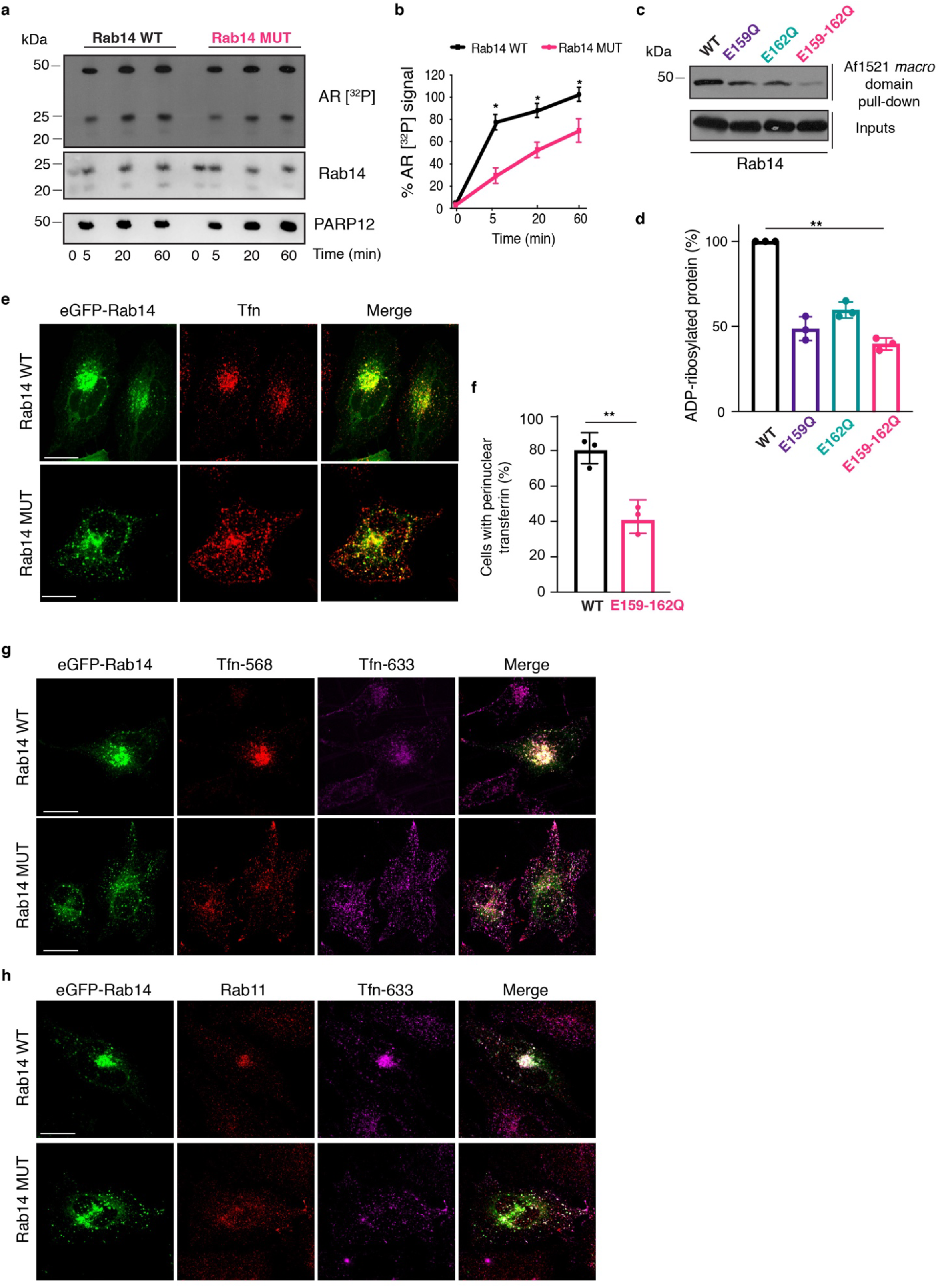
The Rab14-MARylation defective mutant phenocopies PARP12 depletion and PJ34 treatment on transferrin pathway. **(a)** *In vitro* ADP-ribosylation assay using GST-tagged purified PARP12 catalytic fragment and His-tagged purified Rab14 wild-type (Rab14 WT) or its MARylation defective mutant E159Q-E162Q (Rab14 MUT), in presence of 4 μCi of [^32^P]-NAD^+^ and 30 μM of total NAD^+^. Reactions were stopped at different times (as indicated); the incorporated [^32^P]-ADP-ribose was detected by autoradiography (AR [^32^P]). Lower panels show total levels of Rab14 and PARP12. **(b)** Quantifications of *in vitro* MARylated Rab14 are reported in the graph. Data represent the mean ± S.D. (N = 3 independent experiments, two-sided one-sample t-test; *P <0,05). **(c)** Representative Af1521 *macro* domain pull-down assay of total lysates from HeLa cells transfected with EGFP-tagged Rab14 WT (WT) or its MARylation defective mutants (E159Q, E162Q, E159Q-E162Q). Af1521 bound proteins (Af1521 *macro* domain pull-down) and total cell lysates (inputs) were separated by SDS-PAGE and analyzed by western blotting with a Rab14 antibody. **(d)** Quantifications of MARylated Rab14 relative to (c) are reported in the graph. Data represent the mean ± S.D. (N = 3 independent experiments; one way Analysis of Variance; ***P <0,0001). **(e)** Representative confocal microscopy images of transferrin internalization (Alexa-568-Tfn) in HeLa cells transfected with EGFP-tagged Rab14 WT or its MARylation defective mutant E159Q-E162Q (Rab14 MUT). **(f)** Quantification of cells showing internalized transferrin into perinuclear recycling endosomes on the total transferrin fluorescence is reported in the graph. Data represent the mean ± S.D. (N = 3 independent experiments; two-sided one-sample t-test; *P <0,0025). **(g)** Representative confocal microscopy images of double-transferrin internalization in HeLa cells transfected with EGFP-tagged Rab14 WT or its MARylation defective mutant E159Q-E162Q (Rab14 MUT). **(h)** Representative confocal microscopy images of Alexa-633-Transferrin (Tfn) uptake in HeLa cells upon overexpression of EGFP-tagged Rab14 WT or its MARylation defective mutant E159Q-E162Q (Rab14 MUT). Cells were processed for immunofluorescence and Rab11 localization analyzed using a Rab11 antibody (red). Merged signals are also reported. Scale bars, 10 μM.

In parallel, to identify the main substrate site(s) of MARylation, two single point mutants -Mut1 (E159Q) and Mut2 (E162Q)-were also generated and tested for their MARylation in cells. Compared to wild-type Rab14, the mutants binding to Af1521 *macro*-domain was reduced by about 50% (Fig. 5c, d).

The additional two predicted residues (E187, E205) were analysed with a similar strategy. A triple point mutant Q159-Q187-Q205 was generated and tested and did not show any deviation from the 50% reduction carried by E159Q/E162Q (not shown). Thus, we excluded that E187 and E205 are MARylated sites. From here on, the double-mutant Q159-Q162 will be referred to as Rab14-MARylation defective mutant (Rab14 MUT).

In conclusion, our data indicate that the glutamic acids in position 159 and 162 of Rab14 are major MARylation sites for PARP12.

### The Rab14 MARylation-defective mutant impairs the transferrin recycling pathway

The MARylation defective mutant was employed to further characterize the role of Rab14 in the transferrin recycling pathway. Thus, the transferrin-uptake assay (see above and methods) was performed in cells over-expressing Rab14 wild-type or its MARylation defective mutant (Rab14 MUT). As shown above, after 30 min of continuous internalization of Alexa-568 transferrin at 37 °C, Hela cells expressing EGFP-tagged wild-type Rab14 showed a perinuclear accumulation of transferrin-containing endosomes (Fig. 5e; more than 80% of total labelled cells, Fig. 5f). Hela cells expressing the Rab14 MUT showed a 60% reduction of cells with a perinuclear localization of transferrin (Fig. 5f) that was instead accumulated in endosomal structures at the cell periphery (Fig. 5e), a phenotype similar to that observed in cells treated with the PARP-inhibitor PJ34 or PARP12-depleted (Figs. 2, 3).

In line with the results described above, during the transferrin-uptake assay, the Rab14 MUT localized in enlarged vesicles at the cell periphery, co-localizing with RUFY1, a marker of early/sorting endosomes (Suppl. Fig. 6a). To evaluate how MARylation contributes to the different steps of the endocytic pathway, we also analysed the localization of FIP1c/RCP, a known Rab14-Rab11 dual effector, acting downstream of RUFY1^62^. FIP1c/RCP colocalizes with wild-type Rab14 at the endosomes^19^; instead, Rab14 MUT and FIP1c/RCP did not colocalize, indicating that Rab14 MUT was not able to bind FIP1c/RCP and therefore is kept at early/sorting endosomes, preventing the endosomal progression and thus the interaction Rab14-FIP1c/RCP (see model Fig. 7 and Suppl. Fig. 6b).

The defective phenotype associated with the Rab14 MUT was then analyzed with two-consecutive transferrin pulses in Hela cells (as described before) and compared with the phenotype observed for wild-type Rab14. Overexpression of the Rab14 MUT caused an accumulation of both fluorescently-labeled transferrin species at peripheral separated enlarged endosomes, which failed to reach the pericentrosomal ERC (Fig. 5g), in agreement with the results observed when MARylation was impaired by PARP12 depletion (Fig. 4b). These data were confirmed in live-cell time-lapse imaging of transferrin uptake in HeLa cells expressing EGFP-tagged wild-type Rab14 or EGFP-tagged Rab14 MUT (Suppl. Videos 3, 4): while in the wild-type Rab14 expressing cells the bulk of fluorescent transferrin bound to its receptor was targeted to the ERC, the Rab14 MUT cells showed an accumulation of the fluorescent transferrin bound to its receptor in peripheral endosomes, positive also for Rab14 MUT (Suppl. Videos 3, 4).

To summarize, in presence of the Rab14 defective MARylation mutant, endocytosed transferrin rapidly enters the early/sorting endosomes, where it is retained as a consequence of its impeded trafficking through the recycling compartment; under these conditions the Rab14 effector RUFY1 remains on peripheral endosomes (Suppl. Fig. 6), thus preventing the normal membrane flow through the recycling endosome.

FIP1c/RCP is a common effector for Rab14 and Rab11a; the latter is associated to perinuclear recycling endosomes and regulates the last step of the transferrin-recycling pathway^1, 19, 63, 64^;. Of note, in Hela cells expressing Rab14 MUT, also Rab11a shifted from the perinuclear endosomal localization toward a peripheral endosomal localization (Fig. 5h), indicating that a defective MARylation also alters Rab11 localization and thus function^13, 65^.

Overall, our findings demonstrate that MARylation of Rab14 is required for the proper progression of endosomes, thus affecting the entire transferrin recycling pathway.

### Rab14 MARylation is functional to its GTPase activity

Rab proteins function as molecular switches cycling between an inactive GDP-bound and an active GTP-bound form; the latter recruits downstream effectors onto membranes^66^. The localization of Rab proteins is controlled by their nucleotide binding status^67, 68^ which also determines their specificity of interaction; in particular, GTP-bound Rab14 specifically interacts with the effector RUFY1 and in turn with FIP1c/RCP^19, 23^.

On these bases and considering our data showing that Rab14 mutant co-localizes with RUFY1 on sorting endosomes, we investigated whether GTP-loaded Rab14 could be more prone to be MARylated by PARP12. To this end, we performed an *in vitro* ADP-ribosylation assay using purified His-Rab14 protein, previously loaded with either GTPγS or GDP nucleotides (see methods and ^69^). As shown in Fig. 6a, GTP-loaded Rab14 was MARylated more efficiently by PARP12 when compared to GDP-loaded Rab14 (2-fold increase). These data indicate that MARylation is associated to active Rab14. To validate this finding, we generated and purified His-tagged FIP1c/RCP deletion mutant corresponding to the C-terminal region of the protein, specifically the residues 559–649 (His-ΔRCP559-649), which include the RBD domain involved in Rab14 interaction. Then, we used His-ΔFIP1c-RCP 559-649 fusion protein as bait in pull-down experiments to isolate the active Rab14 from cell lysates (as ΔFIP1c-RCP 559-649 was reported to specifically interact with GTP-Rab14^32^), at steady-state or under a transferrin-trafficking pulse (see methods). Fig. 6b shows that, in both conditions, the amount of GTP-Rab14 present in cell lysates was higher upon overexpression of the Rab14 MUT compared to the overexpression of wild-type Rab14, indicating that the GTPase activity is not efficient and as a consequence the pool of GTP-bound Rab14 increases. HeLa cells transfected with Rab14 GTP-locked mutant (Q70L) and GDP-locked mutant (S25N) were used as internal controls of the pull-down efficiency (Fig. 6b).

**Fig. 6:**
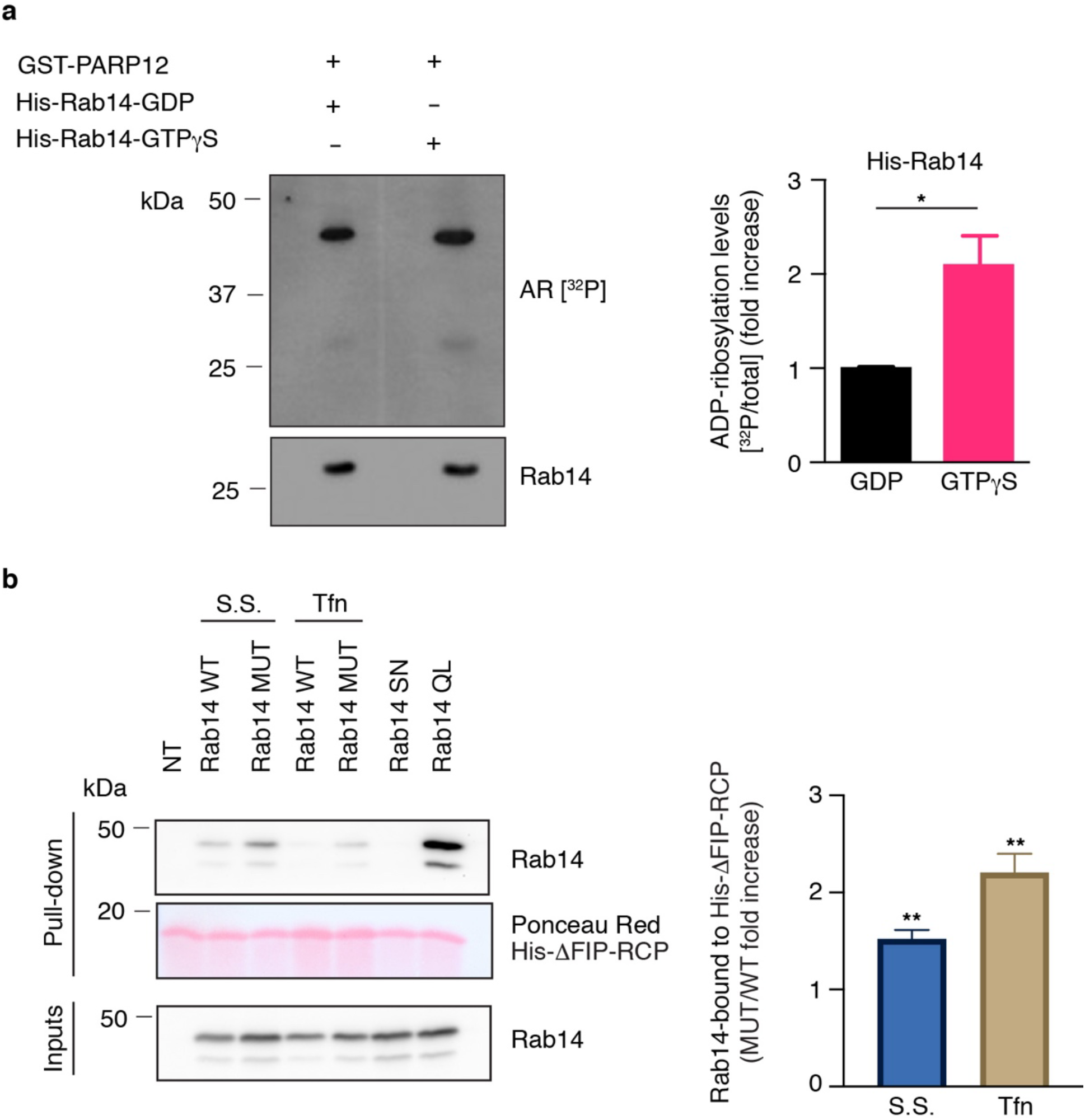
Rab14-MARylation defective mutant is in a GTP-locked state. **(a)** *In vitro* ADP-ribosylation assay using GST-tagged purified PARP12 catalytic fragment and His-tagged purified Rab14, previously loaded with GDP or GTPγS nucleotides, in presence of 4 μCi of [^32^P]-NAD^+^ and 30 μM of total NAD^+^. The incorporated [^32^P]-ADP-ribose was detected by autoradiography (AR [^32^P]). Lower panel shows Rab14 total levels. Quantifications of MARylated Rab14 are reported in the graph. Data represent the mean ± S.D. (N = 3 independent experiments, two-sided one-sample t-test; *P <0,05). **(b)** Representative His-ΔFIP1c-RCP559-649 (ΔFIP1c)-pull-down experiment performed on total lysates of Hela cells transfected with EGFP-tagged Rab14 WT or Rab14 MARylation defective mutant E159Q-E162Q (Rab14-MUT), at steady state (S.S.) or incubated with biotinylated-transferrin (Tfn), as indicated. Total lysates overexpressing Rab14 S25N and Rab14 Q70L were used as negative and positive controls of Rab14 GTP bound state, respectively. ΔFIP1c-bound proteins were eluted and analyzed by western blotting using a Rab14 antibody. Rab14 bound to His-ΔFIP1c indicates the GTP-loaded Rab14 pool. Inputs were also analysed by western blotting using a Rab14 antibody. His-ΔFIP1c protein total levels were detected by Ponceau Red staining. Quantifications of GTP-loaded Rab14 are reported in the graph. Data represent the mean ± S.D. (N = 3 independent experiments, two-sided one-sample t-test; *P <0,05, **P <0,005).

**Fig. 7:**
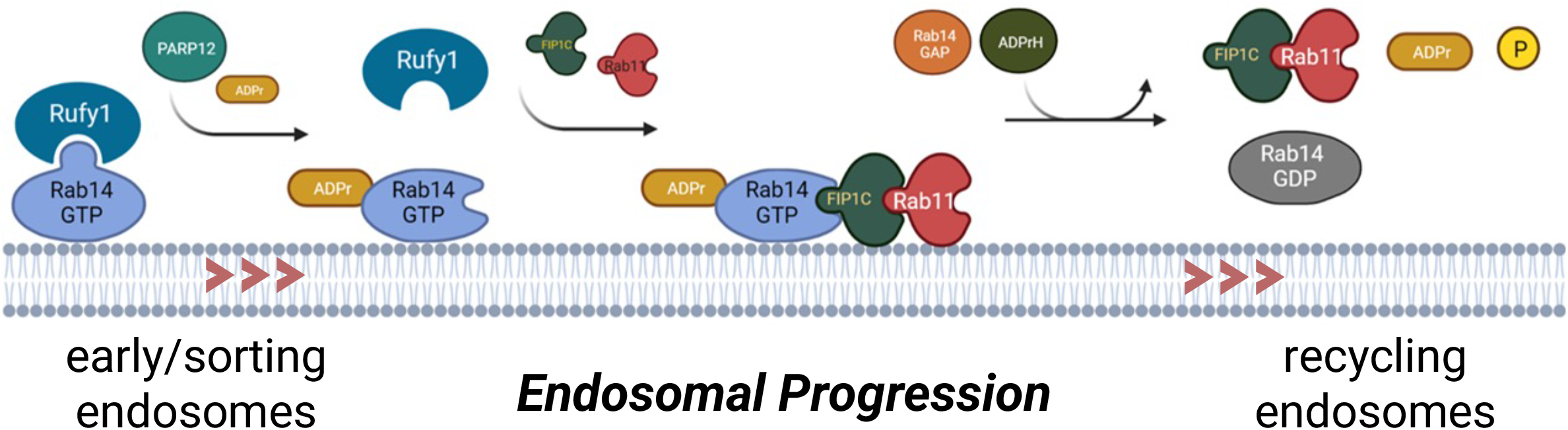
MARylation of Rab14 is required for endosomal progression. Schematic representation of MARylated Rab14 role during endosomal progression. GTP-loaded Rab14 binds the effector RUFY1 on early/sorting endosomes. Endocytic stimuli, such as transferrin bound to its cognate receptor, cause PARP12 translocation to early/sorting endosomes, where Rab14 MARylation occurs. The ADP-ribose likely induces Rab14 conformational changes suitable for binding to the downstream effectors FIP1c/RCP (FIP1C) and Rab11, thus allowing endosome progression towards the endocytic recycling compartment (ERC). Here, GDP-bound Rab14 is released from endosomes, while GTP-loaded Rab11/FIP1C complex allows recycling of endosomes to the plasma membrane. At the ERC, an ADP-ribosylhydrolase (ADPrH) may remove the ADP-ribose moiety, making Rab14 available for the following cycle.

Collectively, these results identify MARylation as a functional PTM needed for the progression of the endosomal compartment toward recycling at the ERC; there, the interaction with the specific effector FIPc/RCP might favor the activation of the Rab14 GTPase activity and conversion to Rab14-GDP needed to maintain an efficient endocytic cycle.

## DISCUSSION

In this study we identify a main event involved in the progression of the endosomes of the endocytic slow-recycling pathway which requires in turn Rab14, Rab11 and the effectors RUFY1 and FIP1c/RCP, whose interactions are determined by a specific modification of Rab14, also affecting its GTP-bound status. This is the MARylation of Rab14 (the first known functional PTM of this small GTPase), that results to be a necessary PTM underlying the endosomal recycling pathway. This modification of Rab14 is due to the mono-ADP-ribosyl-transferase PARP12, an enzyme already involved in the control of the exocytic pathway^46^.

Upon endocytic stimuli, both PARP12 and Rab14 localize at Tfn-positive early/sorting endosomes, where MARylation occurs and allows the progress of the slow-recycling pathway, as demonstrated for the transferrin receptor transport (see model, Fig. 7). These findings, together with our recent report on Golgin-97, point at PARP12 and its substrates as modulators of discrete steps of the exo- and endocytic traffic, and indicate MARylation as a crucial PTM in the control of intracellular membrane transport^45, 46^.

Cellular organization, compartmentalization and cell-to-cell communication are crucially dependent on intracellular membrane trafficking and are accomplished through the coordinated action of highly regulated molecular machineries designated for export, uptake, sorting and recycling of proteins and solutes^1, 6, 70^.

Accordingly, proteins involved in nutrient uptake (such as transferrin receptor for iron, low-density lipoprotein receptor for cholesterol and glucose transporter type 4 for glucose) as well as proteins involved in cell adhesion and migration (such as integrins and cadherins) or proteins involved in cell signalling (such as receptor tyrosine kinases and G protein-coupled receptors) all exploit the endosomal recycling pathway for their correct localization/functions^71^. When defective, this process contributes to the onset and progression of several diseases, including metabolic disorders, cancer, infections as well as some of the most common neurological disorders^72, 73^.

In eukaryotic cells, multiple proteins contribute to the achievement of complex membrane trafficking tasks; in this context, the monomeric small GTPases Rabs play central roles, by binding specific effectors in a coordinated manner and thus establishing inter-connected cascades of events^5, 6, 67^. The molecular determinants controlling this spatio-temporal Rab-membrane recruitment and/or Rab-activation cycle (GDP/GTP cycle) still remain unclear.

PTMs of Rab proteins are emerging as major mechanisms by which this regulation occurs, usually determining Rab binding to specific partners (regulators or effectors;^74, 75^). This is the case of Rab8 phosphorylation, that impairs its GEF-mediated activation (by inhibiting Rab8-Rabin8 interaction^76^) or of Rab7 phosphorylation/dephosphorylation regulating the interaction with its GAP and downstream effector RILP^34, 77^. Similarly, ubiquitination of Rab11a is known to enhance its GTPase activation^78^.

Following on the relevance that the above PTMs have in Rab function, here we present MARylation of Rab14 as a key PTM that controls Rab14 compartmentalization and function at the recycling-endosomal compartment during transferrin uptake; the consequence of Rab14 modification is found in the interaction with its downstream effectors and thus with the Rab14 GTPase cycle.

Accordingly, MARylation of Rab14 serves as modulator of Rab14-RUFY1 interaction and lack of a MARylated Rab14 blocks the canonical flow of the recycling pathway, possibly because of a prolonged interaction of the non-MARylated Rab14 with its specific effector RUFY1.

Of particular interest in this context is a recent study showing that GTP-Rab14 is required for RUFY1 localization at early/sorting endosomes, where RUFY1 binds the dynein/dynactin complex to allow endosome progression along microtubules^56^. These findings support our data showing a key role of MARylated Rab14 in endosomal progression, pointing at a novel PTM central to the process. More specifically, the Rab14 ability to bind both RUFY1, a dual Rab4/Rab14 effector, and Rab11-FIP1c/RCP, a dual Rab14/Rab11 effector, has been interpreted by us as a feature of this regulatory step in the Rab4-Rab14-Rab11 cascade, acting between early/sorting endosomes (characterized by Rab4 and RUFY1) and recycling endosomes (characterized by Rab11 and FIP1c/RCP), along the transferrin receptor slow-recycling pathway^19, 23, 58^.

The pathway we describe herein occurs during transferrin trafficking, when PARP12 can be detected at early/sorting recycling endosomes. We have recently shown that PARP12 localizes at the TGN, where this enzyme controls transport of a subset of basolateral cargoes, through the MARylation of Golgin-97, and that this pathway is part of the PKD signaling activated at the TGN during a traffic wave^46^. Indeed, in both pathways a kinase activity, PKD in exocytosis^46^and tyrosine-kinases in endocytosis^79^ are the trigger for the activation of the transport and possibly of the PARP12 activity, as indicated in the case of the Golgin97 MARylation^46^.

Following on the data reported, we envision a model where, upon transferrin internalization, PARP12 translocates from the TGN to early/sorting endosomes where, by MARylating Rab14, it modulates the Rab14/RUFY1 interaction, possibly releasing RUFY1 while favouring Rab14 subsequent interaction with FIP1c/RCP. The progression of the endosome guarantees Rab14/Rab11-FIP1c/RCP interaction, needed for both the final step of the recycling endosomal process and the control of Rab14 GTPase activity. Whilst RUFY1 cooperates with Rab14 at the sorting endosomes, Rab11-FIP1c/RCP could engage Rab14 on recycling endosomes and promotes its progression towards the ERC^20^ before recycling back to the plasma membrane (see model, Fig. 7).

It is noteworthy that based on our model, the modulation of the Rab14 MARylation by controlling the endosome progression toward the Rab11-specific compartment, could likewise ameliorate all those pathological conditions that have been related to defects in this pathway^24^. This has been the case for neurological disorders such as Parkinson’s or Alzheimer disease where the modulation of the endocytic pathway through Rab11 expression affected α-synuclein aggregation and β-amiloid production^80, 81^. Similarly, several reports have associated Rab14 to cancer development which could be affected by the inhibition of the Rab4-Rab14-Rab11 cascade^26, 32, 82, 83^

While the molecular details of the Rab14, and as a consequence PARP12, activity in the pathogenesis of these diverse diseases have yet to be defined, their mechanistic role in these processes arises the interest in them as druggable enzyme/s amenable to pharmacological development toward therapies.

## METHODS

### Reagents and antibodies

All chemicals and reagents were purchased either from Sigma-Aldrich (Merck) or Thermo Fisher Scientific (Invitrogen). Paraformaldehyde from Electron Microscopy Sciences; [32P]-β-NAD+ (specific activity, 800 Ci mmol−1) from PerkinElmer; Ni-NTA Agarose beads from Qiagen, protein A Sepharose beads from Amersham.

Recombinant protein fragment of human PARP12 was from Abcam (#ab193474).

Commercially available antibodies: mouse monoclonal antibodies to Transferrin Receptor/CD71 (clone H68.4; #13-6890, Thermo Fisher Scientific; western blots (WB) 1:3,000; immunofluorescence (IF) 1:200), EEA1 (#610456, BD Transduction Laboratories; WB 1:1,000; IF 1:100), Golgin-97 (CDF4, #A-21270, Thermo Fisher Scientific; IF 1:100); rabbit polyclonal antibodies to Rab4 (#ab13252, Sigma-Aldrich; IF 1:100), Rab14 (#R0656; Sigma-Aldrich, WB 1:2,000, IF 1:100), Rab11 (#71-5300, Thermo Fisher Scientific; IF 1:50), Rab11-FIP1 (#HPA025960, Atlas Antibodies; WB 1:1,000, IF 1:100), RUFY1 (#PA5-31400, Thermo Fisher Scientific; IF 1:100), PARP12 (#HPA063872, Sigma-Aldrich Prestige Antibodies, WB 1:2000); goat polyclonal antibody to PARP12 (#ab45873, Abcam, IF 1:200); HRP-conjugated secondary antibodies (Calbiochem), HRP-conjugated anti-rabbit secondary antibody, heavy Chain Specific (Sigma-Aldrich); AlexaFluor-488,-568 and -647 conjugated secondary antibodies and AlexaFluor -488, -568, -633 or -647 Conjugate transferrins (Thermo Fisher Scientific, Invitrogen).

### Plasmids

Plasmid vectors encoding for human EGFP-tagged Rab14 wild-type (WT), Rab14S25N and Rab14Q70L mutants were kindly provided by prof. B. Goud (Curie Institute, PSL Research University, Paris); EGFP-Rab14 ADP-ribosylation mutants were generated from EGFP-Rab14 WT. All oligonucleotide primers used for mutagenesis reactions and for molecular biology cloning are listed in Table I and Table II, respectively. All generated constructs were verified by DNA sequencing.

**Table I:**
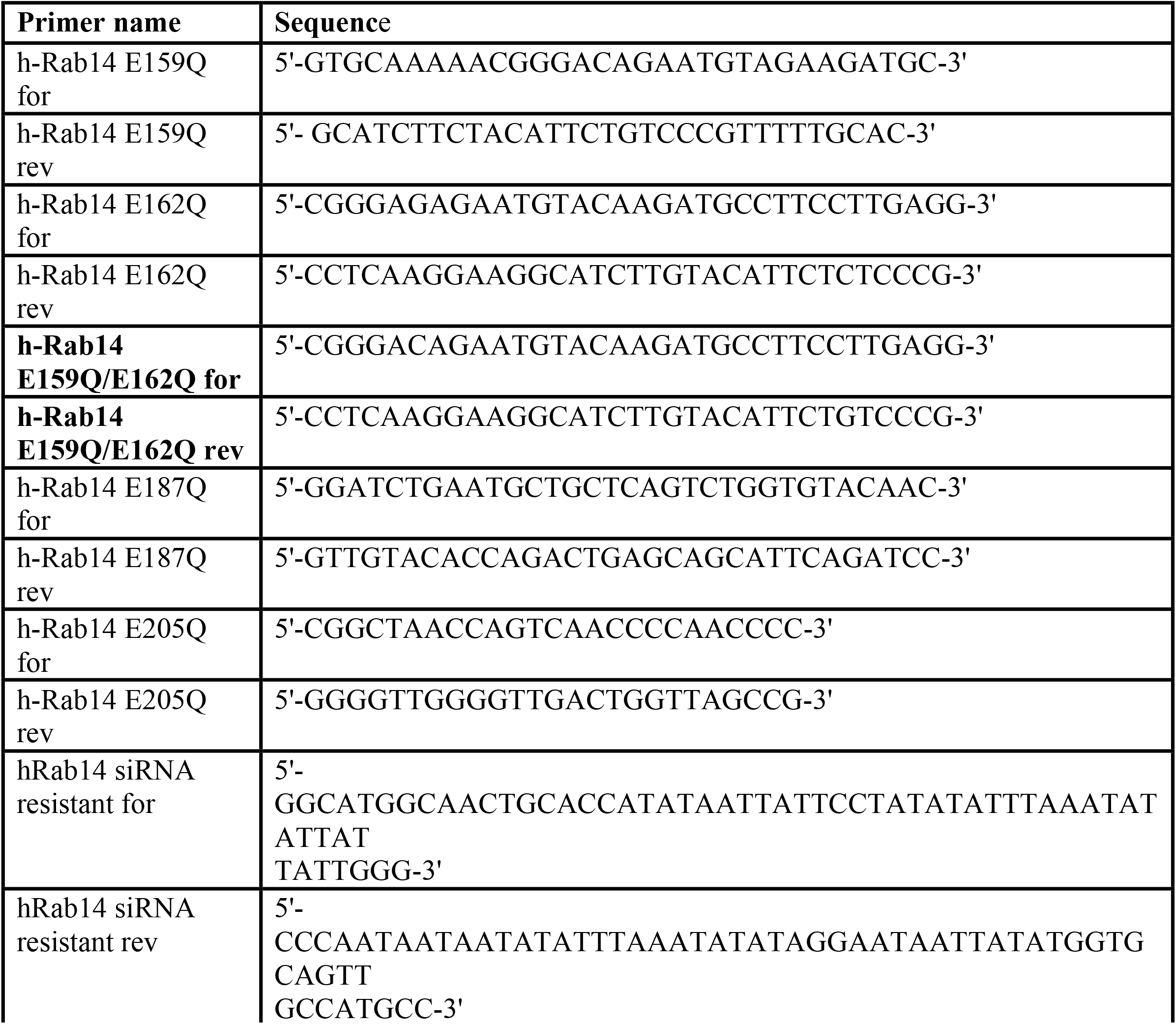
Primers for mutagenesis reactions.

**Table II:**
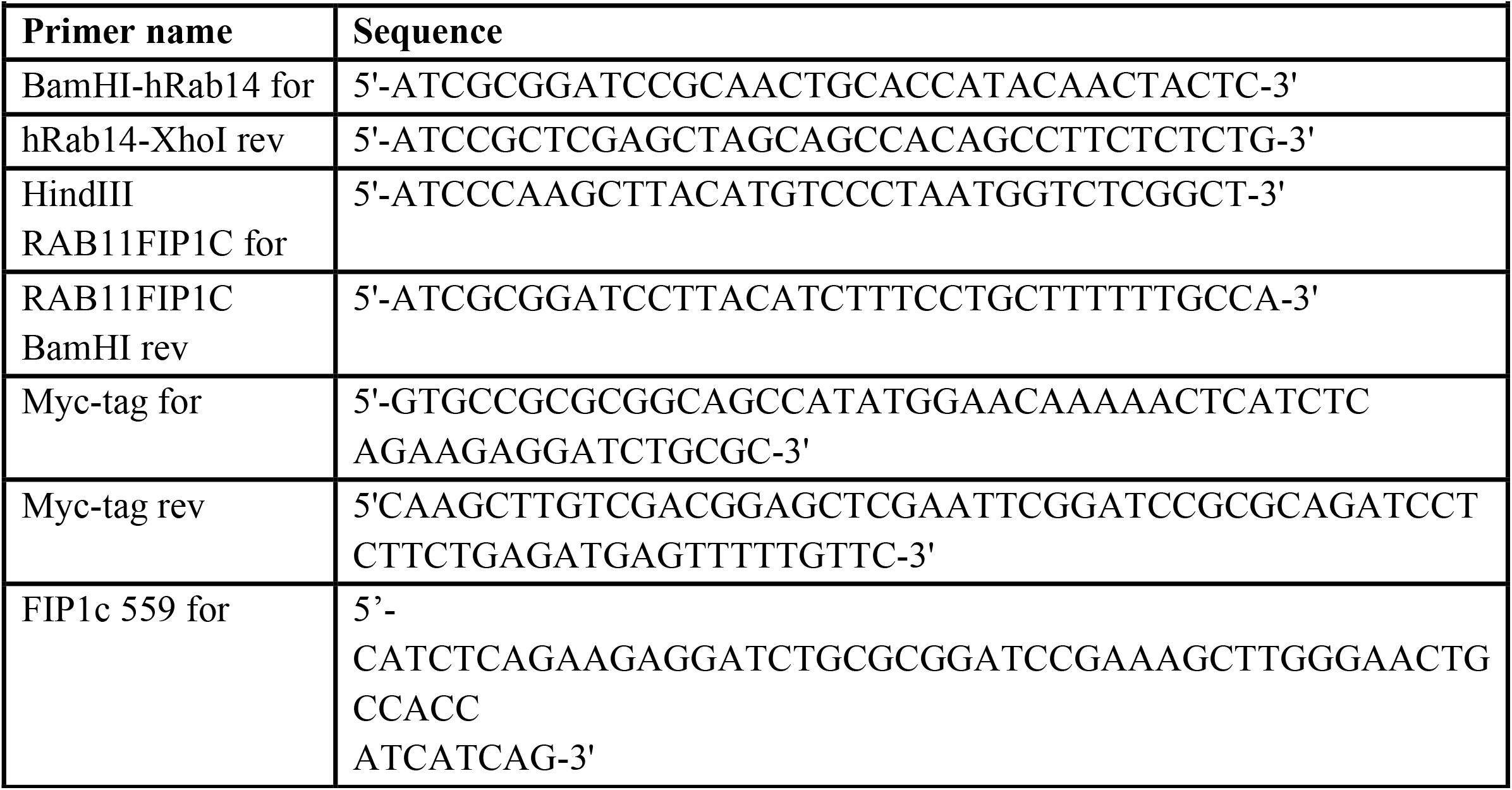
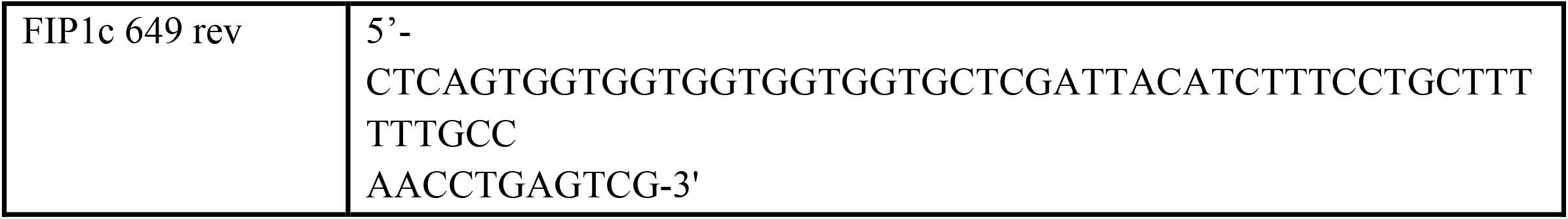
Primers for molecular cloning

#### Generation of human mutant Rab14 constructs

Human Rab14 mutant constructs were generated by site-directed mutagenesis reactions. Human Rab14 E159Q/E162Q double mutant was generated using as template of the mutagenesis reaction the construct encoding the human Rab14 E159Q mutant. Human Rab14 E159Q/E187Q/E205Q triple mutant construct was generated by performing two subsequent mutagenesis reactions: the first one introduced the E187Q mutation into the Rab14 E159Q mutant coding sequence, the second one introduced the E205Q mutation into the Rab14 E159Q/E187Q double mutant coding sequence. The siRNA resistant constructs encoding the human Rab14 WT and mutant proteins were generated by site-directed mutagenesis reactions using as templates the corresponding constructs previously generated. PCR conditions for mutagenesis reactions were as follow: a single denaturation step (95 °C, 5 min) was performed and followed by 20 cycles of denaturation (95 °C, 1 min), annealing (60 °C, 1 min) and elongation (68 °C, 18 min). Then a final elongation step of 10 min at 72 °C was performed. The PCR reaction was carried out in a 50 μl final volume including the following components: 50 ng of cDNA template, 200 μM of each dNTPs, 1x PfuTurbo Cx Hotstart DNA Polymerase buffer, 0.2 μM of each primer and 2.5 U of PfuTurbo Cx Hotstart DNA Polymerase (Agilent Technologies). Following completion of mutagenesis reactions, parental plasmid cDNA was digested by DpnI treatment (20 U/mutagenesis reaction) for 1 h at 37 °C. DNA was then precipitated, resuspended in sterile H_2_O and transformed to chemical competent *E. coli* TOP-10 cells.

The coding sequences of human Rab14 WT and its mutant proteins were cloned into pET-28a(+) prokaryotic expression vector for the expression and purification of the corresponding His N-terminally tagged proteins. Briefly, the coding regions were amplified using BamHI-hRab14 for and hRab14-XhoI rev primers. Then the PCR generated fragments were digested with BamHI and XhoI restriction enzymes and cloned into pET-28a(+) vector, previously digested with BamHI and XhoI restriction enzymes.

#### Generation of human RAB11-FIP1C constructs

Construct encoding the human RAB11 family interacting protein 1 (class I), transcript variant 1, (RAB11-FIP1C) (NM_025151) was purchased from OriGene (#RC209640). Construct encoding the tomato N-terminally tagged human RAB11-FIP1C protein was generated amplifying the human RAB11-FIP1C coding region using HindIII RAB11FIP1C for and RAB11FIP1C BamHI rev primers, digesting the PCR generated fragment with HindIII and BamHI restriction enzymes and finally cloning it into the ptdTomato-C1 vector (Addgene, #54653), previously digested with HindIII and BamHI restriction enzymes.

The construct encoding the His-Myc N-terminally tagged human Rab11-FIP1c C-terminal fragment (559-649 aa) was generated for bacterial expression. The T7-tag region of the bacterial expression vector pET-28a(+) was substituted with a myc-tag by inserting an 86bp fragment by Restriction Free (RF) cloning procedure^84^. The complementary primers used in the RF reaction were Myc-tag for and Myc-tag rev. The C-terminal 559-649aa region of FIP1c was inserted downstream of the myc tag in the modified vector by RF reaction using the FIP1c 559 for and FIP1c 649 rev primers.

### Protein expression and purification

#### His-Rab14 protein purification

BL21 (DE3) pLysS cells transformed with the resulting pET28a plasmids encoding for His-Rab14-WT or His-Rab14-MUT were grown in Luria-Bertani broth, supplemented with 34 μg/ml chloramphenicol and 30 μg/ml Kanamycin, to an OD 600 of approximately 0.6 and induced with 0.5 mM IPTG for 4 h at 37 °C. Bacteria were centrifuged at 6000× *g* for 10 min and resuspended in Lysis Buffer (50 mM Tris–HCl, pH 8.5; 150 mM NaCl; 1 mM DTT) supplemented with protease inhibitors. Then, cells were lysed by sonication on ice and insoluble proteins were removed by centrifugation for 30 min at 10,000× *g* at 4 °C. The supernatant was transferred to a fresh tube and incubated with Ni-NTA agarose resin at 4 °C for 90 min. The beads were then washed three times with Lysis buffer containing 20 mM imidazole and bound recombinant proteins were eluted by using Lysis buffer supplemented with 200 mM imidazole. The elution and collection steps were repeated at least 5 times. The protein peaked in the first two fractions. The fractions containing the greater amounts of protein (at least 0.2 mg/ml) were pooled, dialyzed twice against 1,000 vol. of 20 mM Tris-HCl buffer, pH 8.0, containing 10% glycerol, 0.1M NaCl and 1mM DTT, flash frozen in liquid nitrogen, and stored in aliquots at -80 °C.

#### His-myc-tagged FIP1c C-terminal fragment protein purification (His-ΔFIP1c-RCP 559-649)

BL21 (DE3) cells transformed with the resulting pET28a plasmid encoding for His-FIP1c were grown in Luria-Bertani broth, supplemented with 30 μg/ml Kanamycin, to an OD 600 of approximately 0.6 and induced with 0.2 mM IPTG for 2,5 h at 37 °C. Bacteria were centrifuged at 6000× *g* for 10 min and resuspended in Lysis Buffer (20 mM Tris–HCl, pH 8.0; 300 mM NaCl; 5 mM MgSO_4_; 5 mM DTT; 1 mg/ml Lysozyme) supplemented with protease inhibitors. Then, cells were lysed by sonication on ice and insoluble proteins were removed by centrifugation for 30 min at 10,000× *g* at 4 °C. The supernatant was transferred to a fresh tube and incubated with Ni-NTA agarose resin at 4 °C for 90 min. The beads were then washed three times with Lysis buffer containing 20 mM imidazole and bound recombinant proteins were eluted by using Lysis buffer supplemented with 200 mM imidazole. The elution and collection steps were repeated at least 5 times. The protein peaked in the first two fractions. The fractions containing the greater amounts of protein (at least 0.2 mg/ml) were pooled, dialyzed twice against 1,000 vol. of 20 mM Tris-HCl buffer, pH 8.0, containing, 200 mM NaCl, 1 mM DTT, flash frozen in liquid nitrogen, and stored in aliquots at -80 °C.

### Cell culture, transfections and treatments

HeLa cell lines were from the European Collection of Autenticated Cell Cultures (ECACC, Sigma). Cells were grown in MEM medium supplemented with 10% (vol/vol) FBS, 100 μM MEM Non-Essential Amino Acids Solution, 2 mM L-glutamine, under standard conditions. All cell culture reagents were from Life Technologies. TransIT-LT1 Reagent (Mirus Bio) was used for transient transfection of cDNAs encoding for Rab14 wild-type or its mutants, according to the manufacturer’s instructions. For knock-down experiments, HeLa cells were transfected with 100 nM PARP12 siRNA pool (L-013740-00-0005; Dharmacon); 50 nM Rab14 siRNA (5′-CAACUACUCUUACAUCUUU-3′, Sigma-Aldrich) for 72 h using Lipofectamine 2000, according to manufacturer′s instructions. Where indicated, Hela cells were treated with 50 μM PJ34 (Sigma-Aldrich) for 2 h, 10 μM JW55 (Tocris Bioscience) for 3 h; 10 μM XAV-939 (Tocris Bioscience) for 3 h; 50 μM IWR-1 (Sigma-Aldrich) for 3 h or with 0,1 mM primaquine (Sigma-Aldrich) for 30 min.

### Immunoprecipitations

HeLa cells in 10-cm Petri dishes were transiently transfected with 5 μg of EGFP-tagged constructs, using TransIT-LT1. Twenty hours after transfection, cells were washed three times with ice-cold phosphate-buffered saline (PBS) and lysed in immunoprecipitation buffer (50 mM Tris-HCl, pH 7.5, 150 mM NaCl, 1% TritonX-100 supplemented with Roche protease inhibitor cocktail). Lysates were centrifuged (5,000x *g*, 10 min, 4 °C), and protein concentration determined using Bradford assay. One mg of cell lysate was incubated with 5 μg rabbit anti-Rab14 antibody, at 4 °C on a rotating wheel. After 4 h incubation, 50 μl protein A Sepharose beads were added and incubated for 1 h on a rotating wheel. The beads were washed 6 times with immunoprecipitation buffer. Bound proteins were eluted by boiling the beads in 100 μl Laemmli buffer for 5 min, and then analyzed by SDS-PAGE and western blotting.

### Preparation of total membrane fraction

Confluent HeLa cells plated in 10-cm Petri dishes were washed three times with ice-cold PBS, and mechanically detached in 800 μl HEPES buffer (20 mM HEPES pH 7.4, 1 mM EDTA, 250 mM sucrose) using a cell scraper. The cells were recovered and then sonicated on ice 3 times for 15 s; unbroken cells were removed by centrifugation at 500x *g* for 5 min. The resultant supernatants were ultra-centrifuged for 1 h at 100,000x *g*, with the supernatants representing the cytosolic fraction and the pellets the total membrane fraction. The total membrane fractions were then resuspended in 20 mM HEPES, pH 7.4, containing 1 mM EDTA and protease inhibitors, and stored at -80 °C. A few microliters were lysed in 1 M NaOH and the protein concentration was evaluated by Bradford assay.

### *Macro* domain-based pull-down assays

HeLa cells in 10-cm Petri dishes were lysed in 1 ml RIPA buffer (100 mM Tris HCl, pH 7.5, 150 mM NaCl, 1% Igepal, 0.1% deoxycholate, 0.1% SDS, supplemented with Roche protease inhibitor cocktail), under constant rotation for 30 min at 4 °C. The mixtures were clarified by centrifugation at 13,000× *g* for 10 min at 4 °C. Cell lysates (1 mg) were incubated with 50 μl of a 10 μg/μl GST cross-linked wild-type macro-domain resin at 4 °C, on a rotating wheel. After the incubation, the mixtures were centrifuged at 700× *g* for 5 min to recover proteins bound to the wild-type *macro* domain. The resin was previously equilibrated with RIPA buffer. Resins were then washed 3 times with RIPA buffer and additional 2 times in the same buffer without detergents. Each time, the samples were centrifuged at 700× *g* for 5 min. At the end of the washing steps, the resins were resuspended in 100 μl Laemmli buffer, boiled, analyzed by SDS-PAGE and transferred onto nitrocellulose for western blotting.

### Identification of PARP12 substrates

For the identification of PARP12 substrates by MS analysis, 8 mg total membranes (see above) obtained from HeLa cells transiently transfected with either empty vector or untagged PARP12 (#SC112439, Origene) were used in ADP-ribosylation assays, performed in presence of 2 mM total NAD^+^, at 25 °C for 8 h. At the end of the reaction, membranes were recovered by centrifugation at 100,000x *g* for 60 min at 4 °C. The resulting pellet was solubilized in 4.5 ml RIPA buffer (100 mM Tris HCl, pH 7.5, 150 mM NaCl, 1% Igepal, 0.1% deoxycholate, 0.1% SDS, and protease inhibitors), under constant rotation for 30 min at 4 °C. The mixtures were then clarified by centrifugation at 13,000x *g* for 10 min at 4 °C. Resulting supernatants were first subjected to a pre-clearing step (with 100 μl of a 10 μg/μl GST cross-linked mutated Af1521 *macro* domain resin for 8 h at 4 °C, on a rotating wheel). After this incubation, the mixtures were centrifuged at 500x *g* for 5 min to recover the proteins bound to the mutated Af1521 *macro* domain, and the supernatants were incubated for a further 8 h with the same amount of GST cross-linked wild-type Af1521 *macro* domain resin. The resins were previously equilibrated with RIPA buffer. After 8 h incubation, the samples were centrifuged at 500x *g* for 5 min. Both of the resins were then washed 8 times with RIPA buffer and another 8 times in the same buffer without detergents. Each time, the samples were centrifuged at 500x *g* for 5 min. At the end of the washing steps, the resins were resuspended in 100 μl SDS sample buffer, boiled and analyzed by 10% SDS/PAGE. The gels were then stained with GelCode Blue Stain Reagent. The bands were analyzed by LC-MS/MS (at CEINGE Institute, Naples, Italy). The identified proteins bound to the wild-type Af1521 *macro* domain resins (from both control and PARP12-over-expressing samples) were sub-grouped based on their functions, according to the data available in literature. Only those proteins that were identified in the PARP12 overexpressing sample were considered as PARP12 specific substrates.

### Active Rab14 pull-down

Hela cells plated on 10-cm Petri dishes were transiently transfected with 5 μg of EGFP-tagged Rab14 constructs, using TransIT-LT1, according to the manufacturer instructions. Twenty hours after transfection, cells were washed twice with ice-cold PBS and lysed with 750 μL of ice-cold lysis buffer (25 mM Tris, pH 7.4, 1 mM EDTA, 5 mM MgCl_2_, 1 mM DTT, 0.1 mM EGTA, 100 mM NaCl, 1% Nonidet P-40, 10 mM Imidazole, supplemented with Roche protease inhibitor cocktail). Cell lysates were clarified by centrifugation at 10,000× *g* for 5 min at 4 °C and then incubated at 4 °C under constant rotation with His-ΔFIP1c-RCP559-649 fusion protein, previously bound to Ni-NTA agarose beads. After 30 min incubation, beads were washed 3 times with 1 ml of ice-cold lysis buffer; bound proteins were then eluted with elution buffer (25 mM Tris, pH 7.4, 1 mM EDTA, 5 mM MgCl_2_, 1 mM DTT, 0.1 mM EGTA, 100 mM NaCl, 1% Nonidet P-40, 200 mM Imidazole) for 15 min at 4 °C on a rotating wheel. Inputs (20 μg) and 50% of the eluted proteins were separated by 15% SDS-PAGE. After electrophoresis, samples were transferred to nitrocellulose membrane and immunoblotted with a primary Rab14 antibody.

### ADP-ribosylation assays

Purified recombinant His-Rab14 wild-type or relative mutants (2 μg) were incubated with 500 ng purified PARP12 catalytic fragment in 50 μl ADP-ribosylation buffer [50 mM Tris-HCl pH 7.4, 4 mM DTT, 500 μM MgCl_2_, 30 μM unlabelled NAD^+^ /5 μCi of [^32^P]-NAD^+^] at 37 °C for the indicated times. Reactions were stopped by Laemmli buffer addition and samples were subjected to SDS-PAGE and transferred onto polyvinylidene fluoride membranes (Immobilon-FL, EMD Millipore). The incorporated [^32^P]-ADP-ribose was detected by autoradiography.

Where indicated, His-Rab14 was previously loaded with either GTPγS or GDP, as described by ^69^ Briefly, beads containing immobilized His-Rab14 were washed with nucleotide exchange buffer (NE buffer) containing 20 mM Hepes, 100 mM NaCl, 10 mM EDTA, 5 mM MgCl_2_, 1 mM dithiothroitol (DTT), 1 mM GTPγS, pH 7.5, and incubated for 30 min at room temperature with NE buffer containing 10 mM GTPγS under rotation. Subsequently, buffer was drained out and the wash and incubation procedures described above repeated twice. Then, beads were washed with nucleotide stabilization (NS) buffer containing 20 mM Hepes, 100 mM NaCl, 5 mM MgCl_2_, 1 mM DTT, 1 mM GTPγS, pH 7.5, and further incubated with NS buffer in the presence of 10 mM GTPγS for 20 min at room temperature under rotation. His–Rab14 loaded with GDP was prepared as described above, except for the presence of GDP instead of GTPγS in both NE and NS buffers.

### Detergent-free isolation of cholesterol-rich membrane from trans-Golgi network vesicles

HeLa cells in 10-cm Petri dishes were washed twice with ice-cold PBS, followed by 1 min of incubation in ice-cold osmotic buffer (10 mM Tris-HCl, pH 7.4). Then the osmotic buffer was quickly decanted, and cells were scraped in an ice-cold homogenization buffer (10 mM Tris-HCl pH 7.4, 1 mM EGTA, 0.5 mM EDTA, 0.25 M sucrose, supplemented with Roche protease inhibitor cocktail) and homogenized with ten strokes of a loose-fitting Dounce homogenizer. Cell homogenates were centrifuged for 10 min at 1,000x *g* to obtain post-nuclear supernatants (PNS). The nuclear pellet was resuspended in 1 ml of homogenization buffer, sonicated and stored as the nuclear pellet fraction. PNS supernatants were centrifuged at 8,000x *g* for 20 min at 4 °C. The obtained pellet was then resuspended in 1 ml of homogenization buffer, sonicated by 3 × 5 s bursts with a sonicator at 40% amplitude setting and stored as the p8000 pellet fraction. The post-8,000 *g* supernatant was equilibrated in 100 mM sodium carbonate, pH 11.0 for 5 min and sonicated by 3 × 5 s bursts with a sonicator at 40% amplitude setting. This sonicated fraction was transferred to a 12 ml polycarbonate ultracentrifuge tube (Beckman) and adjusted to 40% (w/v) sucrose (10 mM Tris-HCl, pH 7.4, 1 mM EDTA and 1 mM EGTA) to give a final volume of 4 ml. The sample was overlaid with 4 ml of 35% (w/v) sucrose, followed by 4 ml of 5% (w/v) sucrose, and subjected to discontinuous equilibrium sucrose density gradient centrifugation overnight at 4 °C at 180,000x *g* using a swing-out SW41 Beckman rotor. Then 1 ml gradient fractions were harvested beginning from the top of the tube and subjected to SDS-PAGE and western blotting analysis.

### Continuous Transferrin Uptake Assay in live cell

Hela cells were plated in mat-tek and transiently transfected with EGFP-Rab14 WT or EGFP-Rab14-MUT. At 16 h after transfection, cells were serum starved for 2 h, and then treated or not with 50 μM PJ34 for 30 min before adding labeled transferrin. Then, cells were incubated with Transferrin (Tnf) conjugated to Alexa Fluor 568 (50 μg/ml) in trafficking medium (DMEM, 20 mM Hepes) in presence or not of 50 μM PJ34. Cells were imaged for the duration of the experiment (2 h). A z-stack of five planes was acquired continuously for 2 h (488 and 568 for excitation; PMT: 510–550 nm PMT 560–615 nm; 512×512 pixels; frame average, 4). Movies were generated by compiling three-dimensional maximum-intensity projections with the Zeiss Zen program.

### Continuous Transferrin uptake experiments

Hela cells plated on glass coverslips were either treated with 50 μM PJ34 in serum-free growth medium for 1 h or transfected with a pool of PARP12 siRNAs, for 72 h. Upon the respective treatment times, cells were serum starved for 2 h in serum-free growth medium and then incubated for 30 min with Alexa Fluor -568 (or -633)-conjugated -Transferrin (50 μg/ml). Cells were then washed with an acid solution (0.5 M NaCl-0,5% Acetic acid), washed 5 times with ice-cold PBS and finally processed for immunofluorescence staining (see below).

For the two consecutive pulses of transferrin uptake experiments, Hela cells were first labeled with Alexa Fluor 633-conjugated Transferrin (50 μg/ml) for 1 h at 37 °C, then washed with ice-cold PBS and then labeled with Alexa Fluor 568-conjugated Transferrin (50 μg/ml) for 15 min at 37 °C. At the end of the incubation steps, cells were fixed and processed for immunofluorescence staining (see below).

### Immunofluorescence staining

Cells were fixed with 4% paraformaldehyde for 20 min, washed three times in PBS and then incubated in blocking solution [0.5% (w/v) BSA, 50 mM NH_4_Cl in PBS, pH 7.4, 0.05% saponin] for 30 min at room temperature (RT). Cells were subsequently incubated with the specified antibodies diluted in blocking solution for 2 h at RT or O/N at 4 °C. After incubation with the primary antibody, cells were washed three times in PBS and incubated with the respective secondary antibody diluted in blocking solution. Finally, cells were washed 3 times in PBS and once in sterile water to remove salts. Nuclei were stained with Hoechst 33258. Coverslips were then mounted on glass-microscope slides with Mowiol (20 mg mowiol dissolved in 80 ml PBS, stirred overnight and centrifuged for 30 min at 12,000× *g*). Images were taken using a Zeiss-LSM 700 confocal microscope. Optical confocal sections were taken at 1 Air Unit.

### Identification of ADP-ribosylated residues

The bioinformatic tool *ADPredict* ^61^, available online at www.adpredict.net, was used for the prediction of ADP-ribosylated acidic residues (aspartic and glutamic acids). Residues with the highest score were considered for subsequent mutagenesis analysis.

### Statistics and reproducibility

All of the quantified western-blots and confocal co-localization data are the mean ± SEM of at least three independent experiments. Statistical analyses were performed using Prism 5 (GraphPad Software). *p* values were calculated comparing control and each treated group individually using Student’s t test. All statistical parameters are listed in the corresponding figure legends. For all statistical tests, *P* < 0.05 was considered significant and is indicated by asterisks.

## Acknowledgments

We thank Dr. Carmen Valente (IEOS, Napoli) for help in imaging data acquisition and for reading of the manuscript; dr. Alberto Luini (IESO, Napoli) for insightful discussions, suggestions and critical reading of the manuscript; Dr. D. Spano for initial cloning (IEOS, Napoli); dr. Andrea Rosario Beccari (DOMPE’ Farmaceutici SpA Research Center, L’Aquila) for support on the ADPredict analysis; the BioImaging Facility at the Institute for Endocrinology and Experimental Oncology “G. Salvatore” for support in imaging and data analysis; the Italian Association for Cancer Research (AIRC) (to D.C. IG18776), the PRIN No 20177XJCHX project, the SATIN and CIRO POR-projects 2014-2022 for supporting our work.

## Author Contributions

A.C, G.G. and D.C. designed research; A.C. and L.S. performed research; M.L.M contributed with ADPredict and structural analysis; N.D. contributed with constructs preparation and protein purifications; S.D.P. contributed to imaging data analysis and figure preparation; A.C., G.G. and D.C. analyzed data; and A.C, G.G. and D.C. wrote the paper.

---

**Suppl. Fig. 1:**
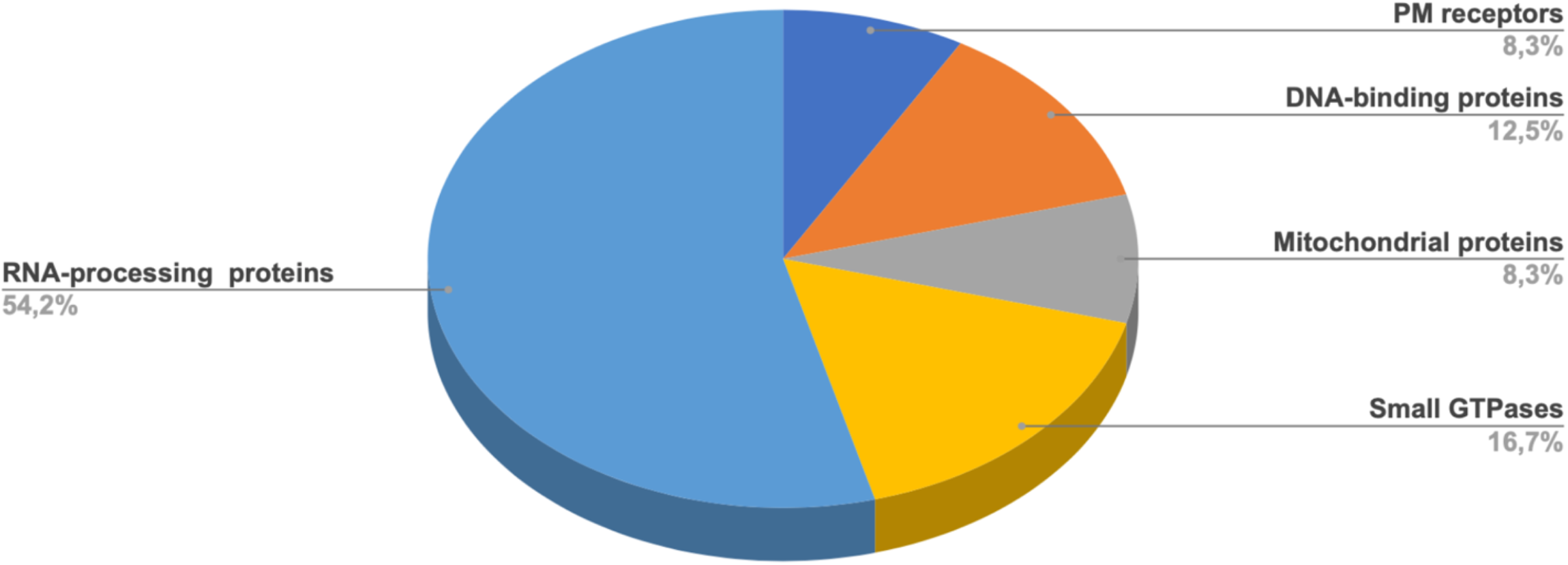
Sub-groups of PARP12 ADP-ribosylated proteins. The substrates identified by MS were clustered into sub-groups based on their functions (as indicated), according to the data available in the literature. Five major sub-groups emerged from this analysis. Each of these is highlighted in a different color.

**Suppl. Fig. 2:**
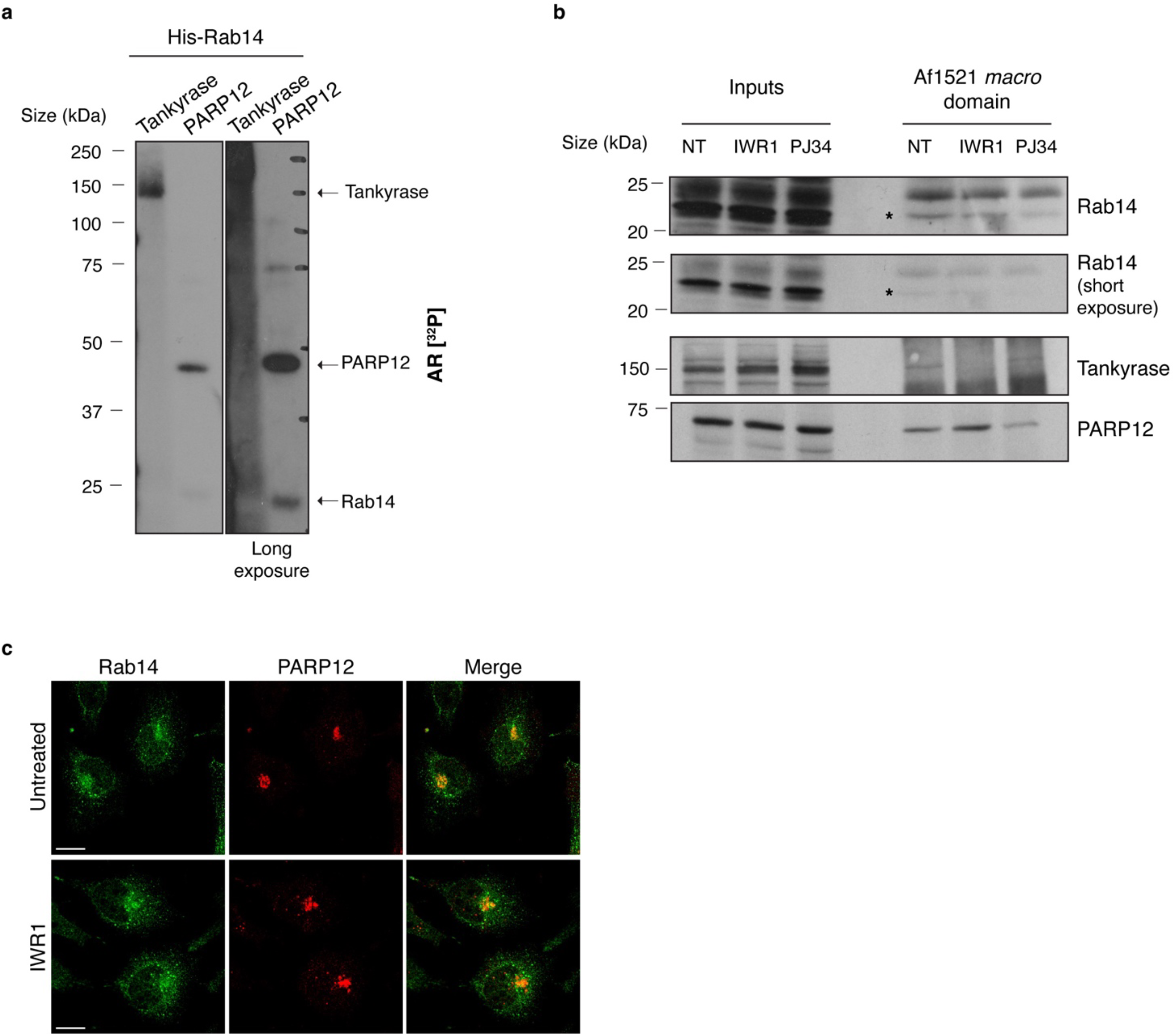
Rab14 ADP-ribosylation is not affected by tankyrase activity. **(a)** *In vitro* ADP-ribosylation assay using purified PARP12 catalytic fragment, tankyrase and His-tagged purified Rab14, in presence of 4 μCi [^32^P]-NAD^+^ and 30 μM total NAD^+^. The incorporated [^32^P]-ADP-ribose was detected by autoradiography (AR [^32^P]). Arrows indicate ADP-ribosylated tankyrase, PARP12 and Rab14. **(b)** Af1521 *macro* domain based pull-down assay of total lysates obtained from (b) HeLa cells treated or not with the broad PARP inhibitor PJ34 (50 μM, 2 h) or (c) with the tankyrase inhibitor IWR1 (10 μM, 2 h) showing MARylated Rab14 (asterisks). ADP-ribosylation of both tankyrase and PARP12 was detected as internal control. **(c)** Representative confocal microscopy images of endogenous Rab14 localization upon IWR1 treatment. After 2 h, cells were fixed and stained with antibodies against Rab14 (green) and PARP12 (red). Merged signals are also shown. Scale bars, 10 μm.

**Suppl. Fig. 3:**
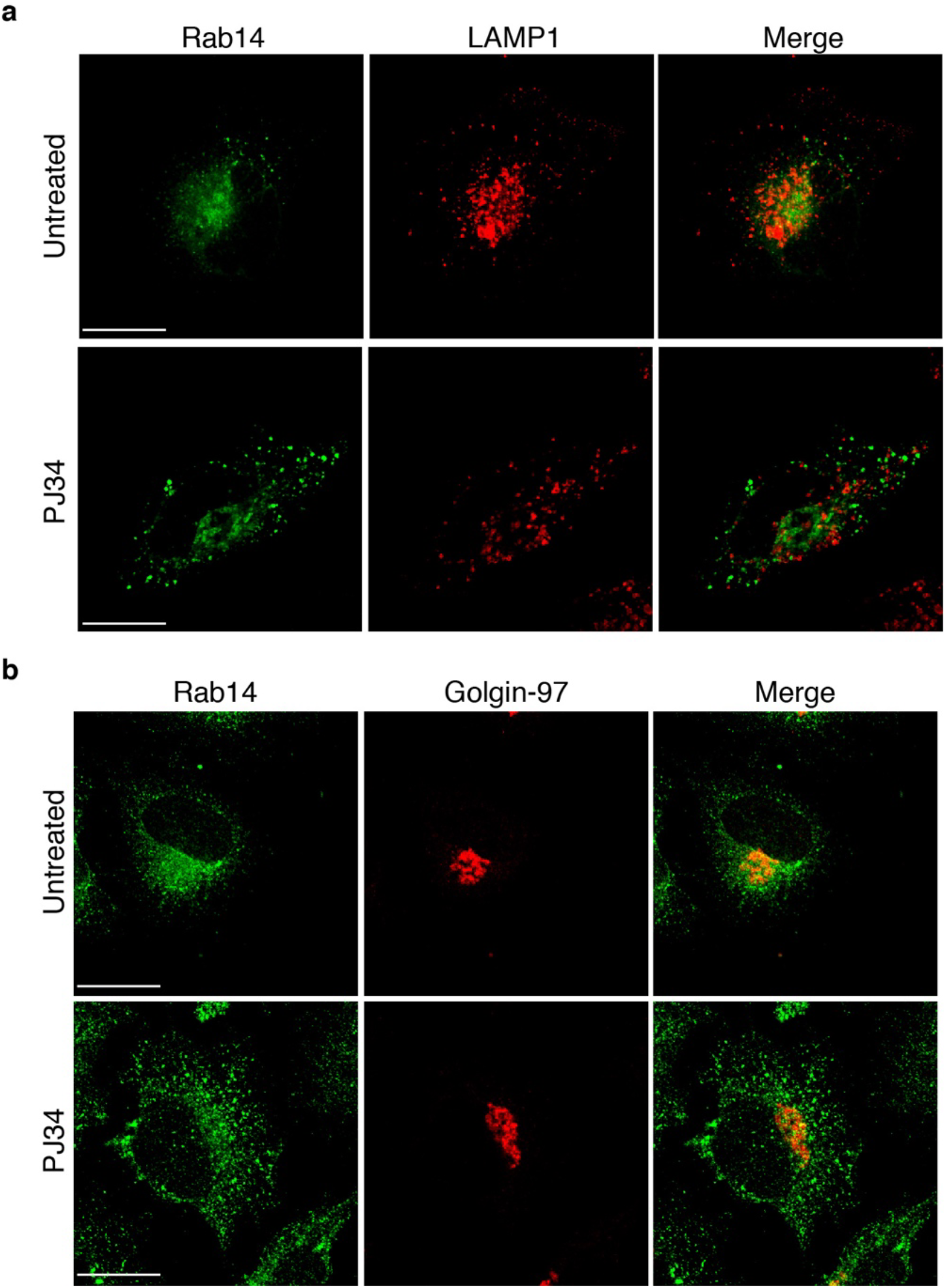
Effect of PJ34 treatment on Rab14 localization. **(a, b)** Representative confocal microscopy images showing endogenous Rab14 localization upon PJ34 treatment (50 μM); after 2 h incubation with the inhibitor, HeLa cells were fixed and labeled with antibodies against (a) Rab14 (green) and LAMP1 (red) or (b) Rab14 (green) and Golgin-97 (red). Merged signals are also shown. Scale bars, 10 μm.

**Suppl. Fig. 4:**
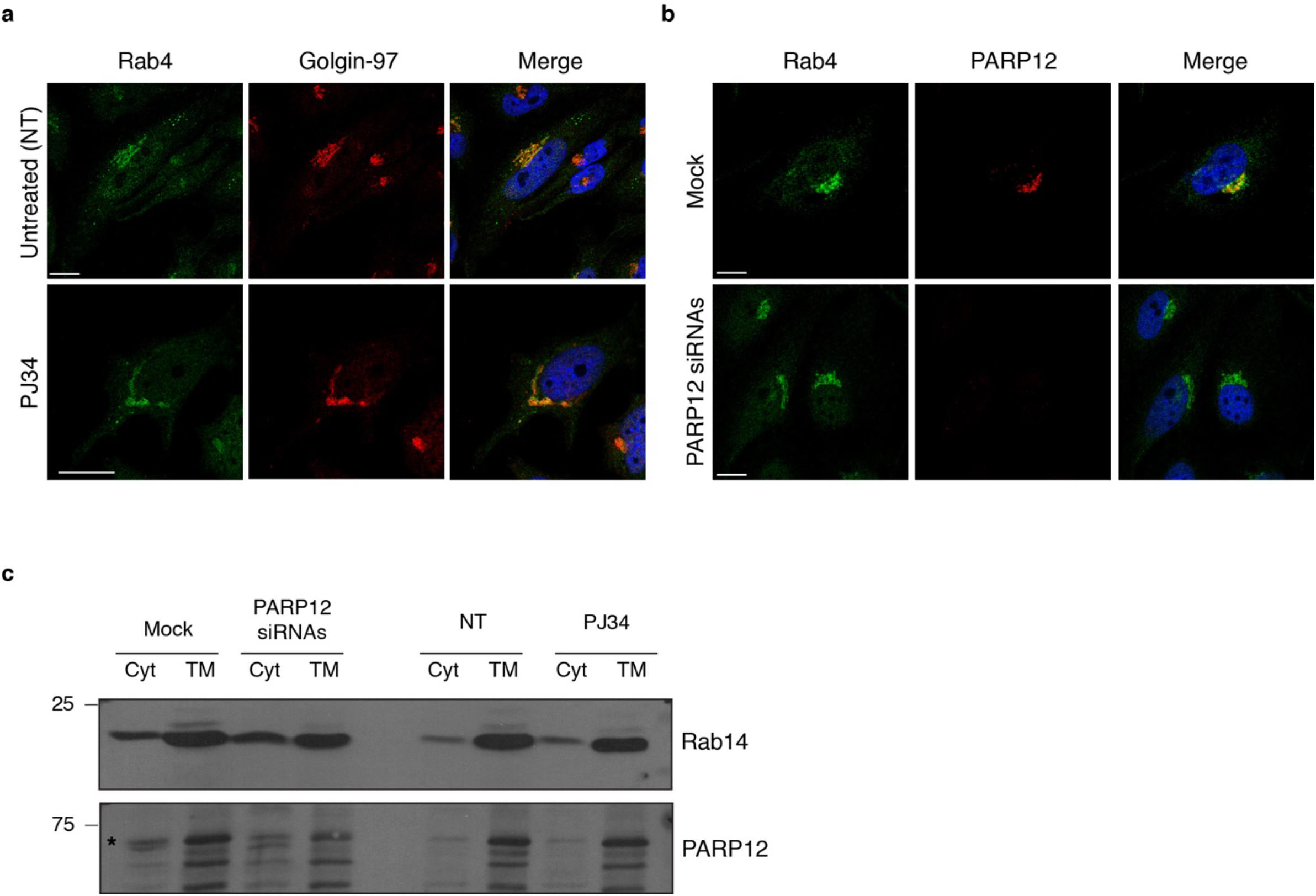
Rab4 localization is not affected by PJ34 treatment or PARP12 knock-down. **(a, b)** Representative confocal microscopy images showing endogenous Rab4 localization upon (a) PJ34 treatment (50 μM, 2 h) or (b) PARP12 transient depletion; after treatments, HeLa cells were fixed and labeled with antibodies against (a) Rab4 (green) and Golgin-97 (red) or (b) Rab14 (green) and PARP12 (red). Merged signals are also shown. Scale bars, 10 μm. **(c)** HeLa cells were transiently transfected with PARP12 siRNAs (72 h) or treated with PJ34 (50 μM, 2 h). Mock and untreated cells were used as controls, respectively. After the specific treatments, cytosolic (Cyt) and total membrane fractions (TM) were prepared and analyzed by western blotting to follow Rab14 distribution. PARP12 was detected as control of the knock-down procedure. Please note that none of the treatments affect the amount of Rab14 in the membrane pool.

**Suppl. Fig. 5:**
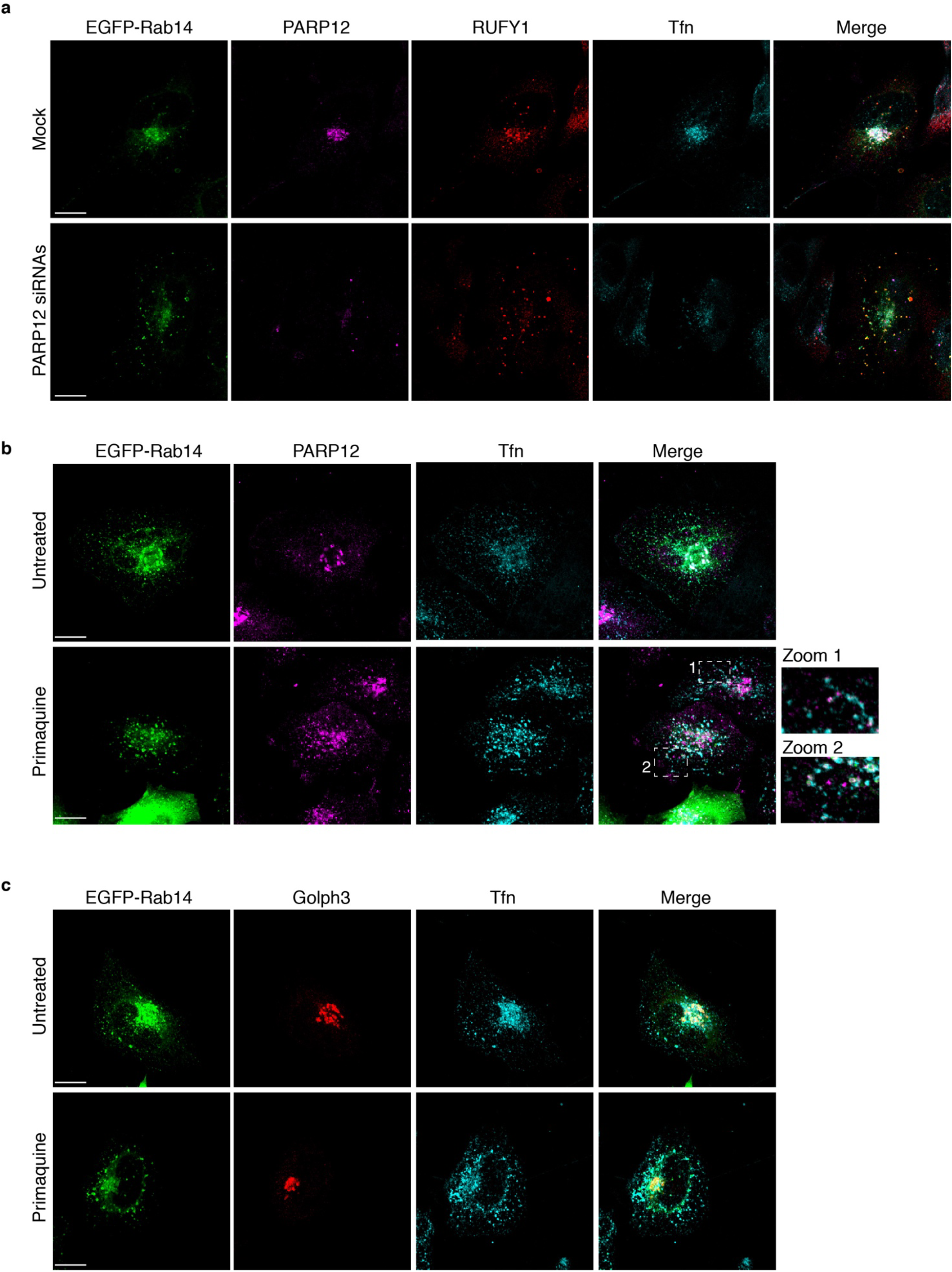
PARP12 depletion blocks transferrin at early/recycling endosomes. **(a)** Representative confocal microscopy images of Alexa-633-Transferrin (Tfn, cyan) uptake in HeLa cells transfected with PARP12 siRNAs and then transfected with EGFP-tagged Rab14. Cells were subjected to a transferrin uptake assay and processed for immunofluorescence with a PARP12 (magenta) and RUFY1 (red) antibodies. **(b, c)** Representative confocal microscopy images of HeLa cells transfected with EGFP-tagged Rab14 and left untreated or treated with Primaquine (0.1 mM), followed by Alexa-647-transferrin uptake (50 μg/ml, 30 min Tfn, cyan). At the end of the assay, cells were fixed and processed for immunofluorescence with (a) PARP12 (magenta) or (b) Golph 3 (red) antibodies. Merged signals are also shown. Scale bars, 10 μm.

**Suppl. Fig. 6:**
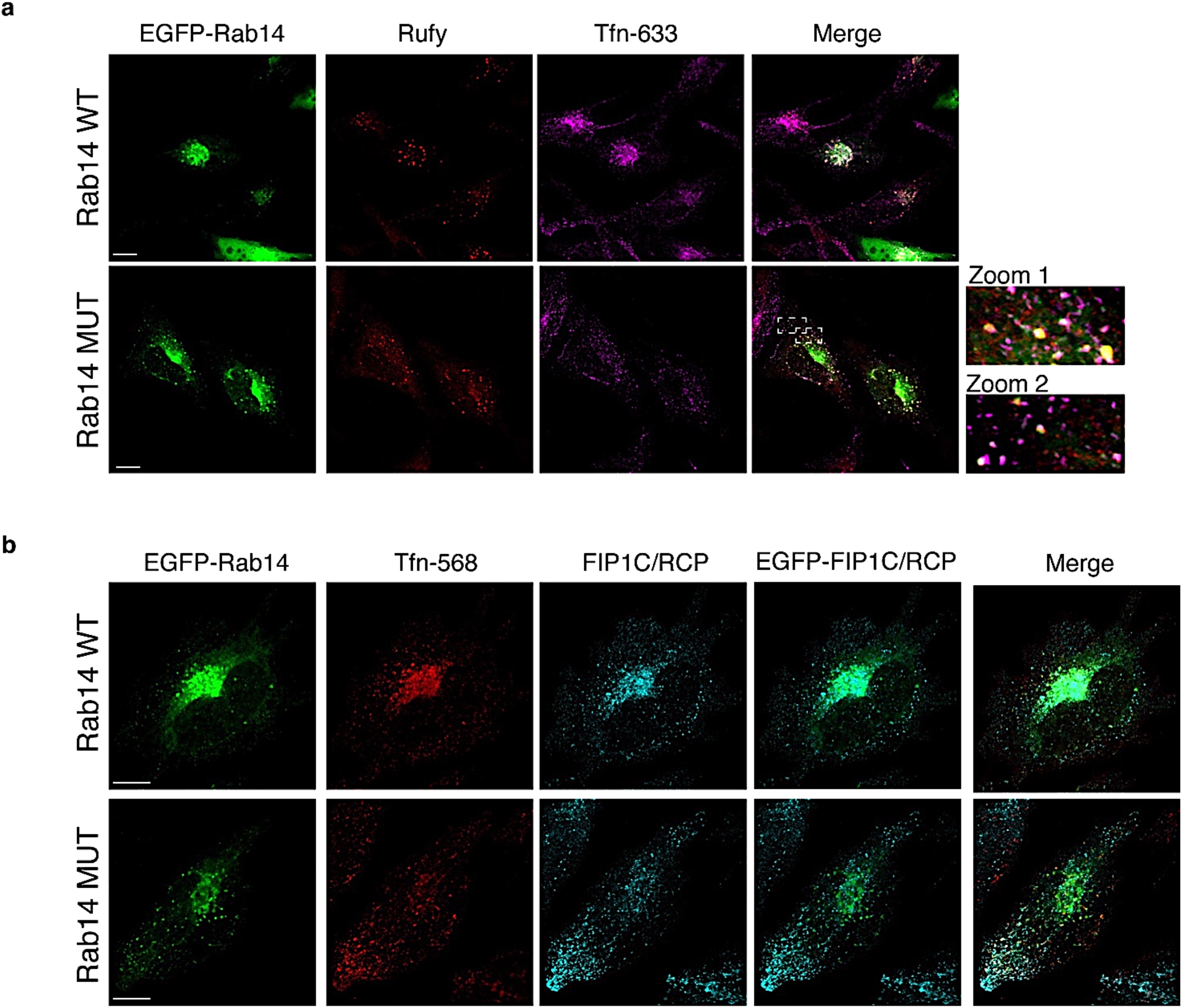
Rab14 MUT blocks transferrin at early/recycling endosomes. Representative confocal microscopy images of transferrin internalization (Alexa-568/633-labeled transferrin, Tnf, as indicated) in HeLa cells transfected with EGFP-tagged Rab14 WT or Rab14 MARylation defective mutant E159Q-E162Q (Rab14 MUT). Cells were processed for immunofluorescence and stained with antibodies against (a) RUFY1 or (b) FIP1c/RCP. Zoom 1 e 2: enlarged view of merged signals. Scale bars 10 μM.

**Table S1:** ADPredict output for human Rab14

| Position | Sequence | Secondary Structure | PDB ID (CHAIN) | ADPredict | AAD RP | AAD RF | AAD SVM | HMD RP | 3DD RP | EXP DATA |
| --- | --- | --- | --- | --- | --- | --- | --- | --- | --- | --- |
| 187 | DLNAAESGVQH |  |  | 0.57 | 0.78 | 0.52 | 0.42 |  |  |  |
| 159 | SAKTGENVEDA | -TTT-TTHHHH | 1Z0F (A) | 0.56 | 0.78 | 0.48 | 0.43 | 0.79 | 0.33 |  |
| 205 | GRLTSEPQPQR |  |  | 0.51 | 0.78 | 0.39 | 0.37 |  |  |  |
| 162 | TGENVEDAFLE | T-TTHHHHHHHH | 1Z0F (A) | 0.45 | 0.78 | 0.17 | 0.17 | 0.46 | 0.67 |  |
| 145 | KQFAEENGLLF | HHHHHHHTT-EE | 1Z0F (A) | 0.38 | 0.67 | 0.24 | 0.22 | 0.11 | 0.68 |  |
| 182 | QDGSLDLNAAE |  |  | 0.38 | 0.60 | 0.31 | 0.21 |  |  |  |
| 127 | IGNKADLEAQR | EEE-TT-GGG- | 1Z0F (A) | 0.37 | 0.60 | 0.14 | 0.18 | 0.59 | 0.33 |  |
| 66 | KLQIWDTAGQE | EEEEEE-TTGG | 1Z0F (A) | 0.36 | 0.60 | 0.10 | 0.20 | 0.59 | 0.33 |  |
| 137 | RDVTYEEAKQF | --S-HHHHHHHH | 1Z0F (A) | 0.36 | 0.27 | 0.20 | 0.08 | 0.59 | 0.68 |  |
| 167 | EDAFLEAAKKI | HHHHHHHHHHHH | 1Z0F (A) | 0.35 | 0.78 | 0.30 | 0.24 | 0.11 | 0.33 |  |
| 129 | NKADLEAQRDV | E-TT-GGG--S | 1Z0F (A) | 0.33 | 0.78 | 0.20 | 0.21 | 0.11 | 0.33 |  |
| 178 | YQNIQDGSLDL |  |  | 0.33 | 0.60 | 0.21 | 0.17 |  |  |  |
| 138 | DVTYEEAKQFA | -S-HHHHHHHH | 1Z0F (A) | 0.32 | 0.60 | 0.07 | 0.09 | 0.59 | 0.23 |  |
| 47 | HTIGVEFGTRI | TS----EEEE | 1Z0F (A) | 0.30 | 0.78 | 0.24 | 0.20 | 0.11 | 0.16 |  |
| 19 | YIIIGDMGVGK | EEEE-STTSSH | 1Z0F (A) | 0.29 | 0.34 | 0.08 | 0.13 | 0.59 | 0.33 |  |
| 39 | KKFMADCPHTI | S---SS-TTS- | 1Z0F (A) | 0.29 | 0.34 | 0.11 | 0.10 | 0.59 | 0.33 |  |
| 163 | GENVEDAFLEA | -TTHHHHHHHH | 1Z0F (A) | 0.29 | 0.35 | 0.07 | 0.24 | 0.46 | 0.33 |  |
| 92 | ALMVYDITRRS | EEEEETT-HH | 1Z0F (A) | 0.27 | 0.26 | 0.12 | 0.07 | 0.59 | 0.33 |  |
| 152 | GLLFLEASAKT | T-EEEE--TTT | 1Z0F (A) | 0.27 | 0.78 | 0.26 | 0.21 | 0.11 | 0.00 |  |
| 108 | SSWLTDARNLT | HHHHHHHHHHHS | 1Z0F (A) | 0.24 | 0.60 | 0.07 | 0.11 | 0.11 | 0.33 |  |
| 144 | AKQFAEENGLL | HHHHHHHHTT-E | 1Z0F (A) | 0.21 | 0.37 | 0.12 | 0.11 | 0.11 | 0.33 |  |
| 33 | LHQFTEKKFMA | HHHHHHS---S | 1Z0F (A) | 0.20 | 0.17 | 0.02 | 0.11 | 0.46 | 0.23 |  |
| 54 | GTRIIEVSGQK | EEEEEEETTEE | 1Z0F (A) | 0.19 | 0.18 | 0.18 | 0.14 | 0.11 | 0.33 |  |
| 71 | DTAGQERFRAV | E-TTGGGT-HH | 1Z0F (A) | 0.19 | 0.12 | 0.12 | 0.04 | 0.46 | 0.23 |  |
| 211 | QPQREGCGC |  |  | 0.18 | 0.37 | 0.11 | 0.07 |  |  |  |
| 133 | LEAQRDVTYEE | -GGG--S-HHH | 1Z0F (A) | 0.11 | 0.26 | 0.08 | 0.12 | 0.11 | 0.00 |  |

